# Sequential exposure to anoxic/oxic conditions leads to biotransformation and detoxification of sitagliptin in urban hyporheic zones

**DOI:** 10.1101/2025.06.03.657625

**Authors:** Simon Klaes, Kerstin Gerundt, Darja Deobald, Luise Henneberger, Beate Escher, Lorenz Adrian, Myriel Cooper

## Abstract

Pharmaceuticals are increasingly recognized as contaminants of concern in aquatic environments. Sitagliptin, an antidiabetic drug that carries a trifluoromethyl group, which is a precursor of the persistent trifluoroacetic acid, is excreted largely unmetabolized and inefficiently removed in wastewater treatment plants, leading to its widespread detection in surface waters. The hyporheic zone — a region between surface water and groundwater — serves as a natural bioreactor with high microbial activity and diverse redox conditions, offering the potential for sitagliptin attenuation. This study explored the biotransformation of sitagliptin in hyporheic sediments under varying redox conditions through batch experiments and field observations. Furthermore, we showed that batch experiments can complement field observations to capture both mechanistic insights and their environmental relevance. Batch experiments revealed amide hydrolysis and N-acetylation of sitagliptin under anoxic conditions, with subsequent deamination and oxidation of transformation products under oxic conditions. Metagenome-resolved metaproteomics suggested *Pseudomonas asiatica* as a key player in the oxic transformation. Field analysis of pore water samples identified up to 6.47 µg L⁻¹ sitagliptin and ten transformation products with concentrations of up to 4.82 µg L⁻¹. Amide hydrolysis products were the most abundant transformation products and preferentially formed under anoxic conditions. All investigated transformation products exhibited lower cytotoxicity and oxidative stress response than sitagliptin in *in vitro* bioassays, highlighting the detoxification potential of the hyporheic zone. By identifying conditions that promote sitagliptin transformation and characterizing its transformation products toxicologically, our work provides parameters for enhanced sitagliptin removal in aquatic environments and improved risk assessment of fluorinated trace organic contaminants.

**Graphical abstract:** 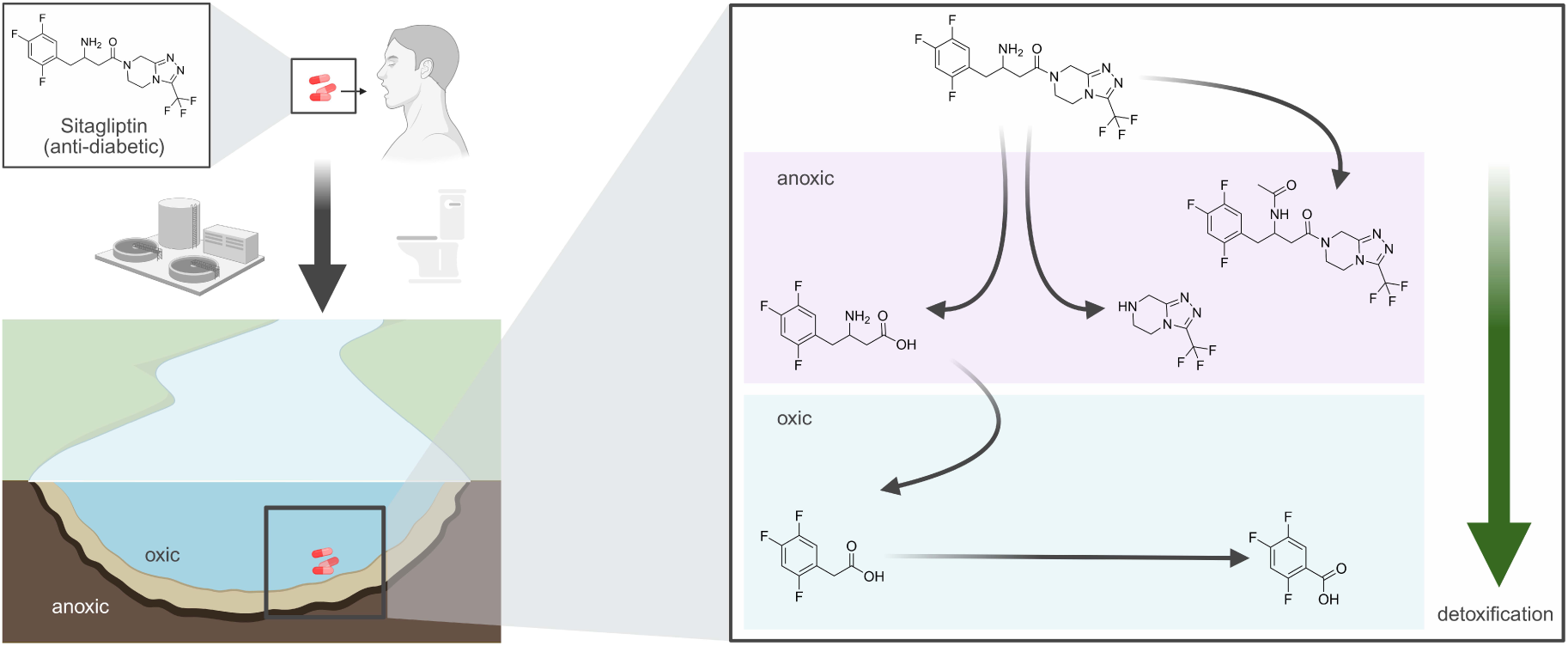

## Introduction

Trace organic compounds (TrOCs), including per- and polyfluoroalkyl substances (PFAS), have become ubiquitous in aquatic systems and pose a growing environmental and public health concern [1–4]. In PFAS, hydrogen substituents are either completely or partially replaced by fluorine substituents, improving the molecule’s thermal and chemical stability. Therefore, PFAS often exhibit exceptional persistence toward abiotic and biotic removal processes and may accumulate in the environment [5].

Sitagliptin (C_16_H_15_F_6_N_5_O), an antidiabetic prescribed in over 130 countries, is one such TrOC and PFAS containing a polyfluorinated aromatic ring system and a trifluoromethyl (C-CF_3_)-functional group, which is a precursor of the very persistent and very mobile trifluoroacetic acid (TFA) [6–8]. Since 2015, sitagliptin has consistently ranked among the top 100 most prescribed drugs in the US, the largest pharmaceutical market globally [6, 9]. In the human body, sitagliptin is only marginally metabolized, with approximately 80% of the dose excreted unchanged via urine, leading to its transport to municipal wastewater treatment plants (WWTPs) [10].

The removal of sitagliptin in WWTPs ranges from approximately 3% to 61% [11–13], with one study reporting no observable degradation in activated sludge treatment [14]. The primarily observed attenuation processes in WWTPs are transformation reactions, including amide hydrolysis and conjugation [11]. As a result, WWTPs discharge sitagliptin and its transformation products to the environment where they may accumulate, particularly in (partially) closed water cycles, such as in Berlin, Germany [15, 16]. In multiple countries, sitagliptin has been detected in WWTP discharge at low µg L⁻¹ concentrations and in river water at high ng L⁻¹ concentrations [11, 13, 17–19]. Exceptionally high sitagliptin concentrations of up to 6.8 µg L⁻¹ were reported in Berlin’s River Erpe, which consists of up to 80% of WWTP discharge [20–22]. Once discharged into the environment, TrOCs such as sitagliptin can undergo various attenuation processes, including dilution, photodegradation, sorption, other abiotic reactions, and biotransformation [4]. According to OECD studies, sitagliptin is stable against hydrolysis and photolysis, and shows only slight biodegradability, but can be adsorbed to sediment in aquatic environments [23].

The hyporheic zone of receiving streams has gained increasing attention as a potential post-WWTP compartment for TrOC attenuation [24]. The hyporheic zone is beneath the stream of a river where surface water and groundwater mix and represents a distinctive habitat [25]. The hyporheic zone is characterized by high microbial activity and redox zonation, providing diverse conditions for transformation reactions. Therefore, the hyporheic zone is often referred to as a “natural bioreactor” or a “river’s liver” [26, 27]. Some TrOCs are more efficiently transformed by microbial communities in hyporheic zones than in WWTPs, highlighting the unique metabolic capabilities of microbial communities in the hyporheic zone [28, 29]. Laboratory [30, 31] and field [20, 32] studies suggest that the hyporheic zone also contributes to the attenuation of sitagliptin. However, the underlying transformation processes, the resulting transformation products, and the ecotoxicological implications of the transformation processes remain unclear.

Sitagliptin acts as a dipeptidyl peptidase-4 inhibitor, showing limited toxicity at environmentally and therapeutically relevant concentrations [10]. Ecotoxicological studies reveal no adverse effects on green algae (72-hour exposure, < 2.2 mg L⁻¹), *Daphnia magna* (21-day exposure, 9.8 mg L⁻¹), and *Pimephales promelas* (33-day exposure, 9.2 mg L⁻¹) [23]. Clinical safety data from a large cohort study (n=6139) demonstrate no adverse effects on humans at therapeutic doses (100 mg/day) over two years [33], although a smaller study (n=104) suggests potential cytotoxic and genotoxic effects at the same dose [34]. *In vitro* studies indicate cytotoxic and genotoxic effects at high concentrations of 200 to 1000 mg L⁻¹ corresponding to 0.49–2.46 mM [35–37]. Toxicological data on transformation products (TPs) of sitagliptin are limited to a few TPs but indicate no increase in toxicity [38, 39]. While TPs often exhibit reduced toxicity, hydrophobicity, and persistence compared to parent compounds, some demonstrate increased toxicity or environmental persistence [40–42]. Thus, TPs need to be included in risk assessment [43].

Here, we investigated biotransformation of sitagliptin with inocula from the hyporheic zone through batch experiments under different redox conditions. To connect the batch experiments with the field, we screened pore water samples from the hyporheic zone across a depth profile at two different sampling sites with contrasting sediment compositions. In addition, we assessed the ecotoxicological impact of sitagliptin and its TPs using *in vitro* bioassays for cytotoxicity, genotoxicity, and oxidative stress response. Our findings contribute to an improved understanding of sitagliptin transformation in the hyporheic zone. With the identification and toxicological description of sitagliptin TPs, our work enables enhanced monitoring and risk assessment of fluorinated TrOCs in aquatic environments.

## Material and Methods

### Chemicals

Sitagliptin was purchased from Biosynth (Compton, United Kingdom). Diphenhydramine hydrochloride and sulfamethoxazole-D4 (SMX-D4) were obtained from Sigma-Aldrich (Taufkirchen, Germany). TP449 (N-acetyl sitagliptin) was purchased from Enamine (Frankfurt (Main), Germany). TP192 (3-(trifluoromethyl)-5,6,7,8-tetrahydro-[1,2,4]triazolo[4,3-a]pyrazine) was supplied by FluoroChem (Hadfield, United Kingdom). TP406 (1-(3-(trifluoromethyl)-5,6-dihydro-[1,2,4]triazolo[4,3-a]pyrazin-7(8H)-yl)-4-(2,4,5-trifluorophenyl)butane-1,3-dione), TP233 ((R)-3-amino-4-(2,4,5-trifluorophenyl)butanoic acid), TP190 (2,4,5-trifluorophenylacetic acid), and TP176 (2,4,5-trifluorobenzoic acid) were purchased from BLD Pharmatech (Reinbek, Germany). All chemical standards were of > 95% purity grade. Formic acid for LC-MS/MS was purchased from SERVA (Heidelberg, Germany), and ultrapure water was prepared with a Merck Milli-Q system (Darmstadt, Germany).

### Sampling site

Sediment cores and pore water were sampled from the River Erpe, a lowland River southeast of Berlin, Germany. The River Erpe is dominated by effluent from the wastewater treatment plant (WWTP) Münchehofe leading to high concentrations of micropollutants in the µg L⁻¹ range [21]. Sediment cores were taken at the Heidemühle site (52.478639 N, 13.635139 E), located approximately 1 km downstream of the discharge point of the WWTP Münchehofe, on April 2021 (and stored at 4 °C before inoculation), September 2022 (directly used for inoculation), and July 2024 (directly used for TP192 and TP233 experiments) [44]. The sediment of this site has previously been characterized by others [44]. For sampling, polyvinyl chloride (PVC) pipes (length 60 cm, inner diameter 6 cm, wall thickness 3 mm) were pushed into the sediment, withdrawn, and sealed with butyl rubber stoppers before transporting them to the laboratory. Sediment was removed as inoculum from the column with a sterilized spatula via pre-drilled ports at specific depths aiming at enriching organisms growing under the corresponding redox conditions based on previous studies of the same sample site [44]. Sediment was taken from 0; 5; 16; and 22 cm sediment depth for oxic, nitrate-reducing, sulfate-reducing, and methanogenic setups, respectively.

Porewater samples were taken with multi-level samplers at two different sites of the River Erpe. Site A (muddy sediment, 52.480237 N, 13.636874 E) was sampled on August 22, 2023 and Site B (sandy sediment, located approximately 150 m downstream of Site A, 52.478985 N, 13.635809 E) was sampled two weeks later on September 06, 2023. For this purpose, multi-level samplers were installed one week prior to sampling for equilibration. Surface water and pore water at sediment depths of 5; 15; 25; 40; and 60 cm were sampled using an IPC 24 multi-channel peristaltic pump (Ismatec, Glattbrugg, Switzerland) with a pumping rate of 5 ml/min. Samples were immediately filtered through 0.45 µm regenerated cellulose syringe filters into different vessels. Samples for Fe²⁺ quantification were acidified to pH 2 by adding 2M HNO_3_. All samples were transported to the laboratory, and stored at 4°C until analysis.

### Cultivation

Sediment enrichment cultures were set up under oxic and anoxic (nitrate-reducing, sulfate-reducing, and methanogenic) conditions. The medium was composed of 200 mg L⁻¹ KH_2_PO_4_, 270 mg L⁻¹ NH_4_Cl, 1000 mg L⁻¹ NaCl, 410 mg L⁻¹ MgCl_2_·6H_2_O, 520 mg L⁻¹ KCl, and 150 mg L⁻¹ CaCl_2_·2H_2_O and was supplemented with 0.1% (v/v) trace element solution SL-10 and 0.1% (v/v) Se/W solution [45]. 1 mM K_2_SO_4_ and 1 mM KNO_3_ were added to sulfate- and nitrate-reducing setups, respectively. 0.1% (w/v) Na-resazurin was added to all anaerobic setups before sparging the medium with N_2_ for 60 min. The sparged medium was aliquoted into serum bottles inside an anaerobic tent (Coy Laboratory Inc., USA) containing a gas atmosphere of 97% N_2_ and 3% H_2_. Serum bottles were crimped with butyl rubber stoppers and autoclaved. After autoclaving, anaerobic setups were amended with 0.1% (v/v) of a vitamin solution [46], 10 mM NaHCO_3_ as buffer, 4 mM L-cysteine as reducing agent (except for nitrate-reducing conditions). Oxic setups were supplemented after autoclaving with 0.1% (v/v) of a vitamin solution [46] and 10 mM HEPES, pH 7.5, as a buffer. Finally, setups were inoculated with 1% (w/v) hyporheic sediment in an anaerobic tent (anaerobic setups) or laminar flow hood (aerobic setups) and spiked with 700 nM sitagliptin from an acetonitrile stock solution (final acetonitrile concentration: 5.7 mM). Sulfate-reducing and methanogenic setups were supplemented with 17 mM H_2_ (nominal concentration) as a putative electron donor. For specific TPs identified in anoxic setups, oxic setups were prepared to test for further transformation. These oxic setups were spiked with 10 nM to 10 µM of TP 193 and TP 233, respectively. All setups were prepared in triplicate and incubated in the dark at 20 °C, with oxic setups continuously shaken and anaerobic setups remaining static. Samples were taken from oxic setups with a pipette and from anoxic setups with a sterile syringe under a laminar flow hood. Anoxic setups were pressurized after sampling to contain about 17 mM H_2_ again.

### Abiotic pH stability test

To test for abiotic sitagliptin transformation at various pH values, the same medium composition as for the methanogenic setup was used but NaHCO_3_ was omitted. Instead, the pH was adjusted to pH 7; 8; 9; or 10 by adding NaOH. Sitagliptin was added to the setups, resulting in a final concentration of 50 µM. The setups were incubated anoxically in the dark. Samples were taken aseptically with a sterile syringe, filtered using 0.2 µM PTFE-HI filters (Agilent, Germany), and sitagliptin and TPs were quantified via LC-MS/MS.

### Quantification of sitagliptin and transformation products via LC-MS/MS

Sitagliptin and TPs were quantified with a liquid chromatography system coupled to tandem mass spectrometry (LC-MS/MS) using a targeted multiple reaction monitoring (MRM) approach. Prior to injection, samples were filtered by 0.2 µM PTFE-HI filters (Agilent, Germany), diluted, and supplemented with an internal standard (diphenhydramine for microcosm experiments and SMX-D4 for porewater samples). The analysis system was composed of a 1260 Infinity II Series LC (Agilent, Germany) and a QTRAP 6500+ MS/MS (AB Sciex, Germany) equipped with a Turbo V ion source switching between positive and negative ionization modes within measurements. Chromatographic separation was achieved at 35 °C on a Zorbax Eclipse Plus Rapid Resolution HT-C_18_ column (100 mm × 3.0 mm, 1.8 μm particle size, Agilent, Germany) equipped with a C_18_ SecurityGuard cartridge (4 mm x 2 mm, Phenomenex, Germany). Water spiked with 0.2% formic acid (A) and methanol (B) was used in a gradient program (Supplementary Table 1) as eluents with a flow rate of 0.4 ml min^-1^. MRM transitions were optimized by direct injection of reference standards or literature references (Supplementary Table 2). Data were analyzed using Analyst 1.7.2 (AB Sciex, Germany).

Details on other analytical techniques can be found in the Supplementary Material.

### DNA analysis

DNA was extracted from frozen cell pellets of biological triplicates using the DNeasy PowerSoil Pro kit (Qiagen, Hilden, Germany) following the manufacturer’s instructions, and stored at -80 °C. Extracted DNA was sent to BGI Tech Solutions (Hong Kong) for 16S rRNA gene amplicon and metagenome sequencing. For 16S rRNA gene amplicon sequencing, the V3–V4 hypervariable region was amplified using the 338F (ACTCCTACGGGAGGCAGCAG) and 806R (GGACTACHVGGGTWTCTAAT) primers and sequenced on the DNBSEQ-G400 platform in the paired-end mode with a read length of 300 bp per read. Raw data were filtered by BGI as follows: 1) primer and adapter sequences were truncated with cutadapt [47]; 2) reads with average Phred quality values< 20 across a 30 bp sliding window were truncated with removing reads whose length was reduced to < 75% of the initial length; 3) reads with ambiguous bases or low complexity (10 consecutive same base) were removed. The filtered data were analyzed using DADA2 v1.34.0 [48] on the Galaxy [49] instance of the Helmholtz Centre for Environmental Research – UFZ using default settings. In brief, forward and reverse reads were merged, requiring no mismatches and a minimum overlap of 12 bp. Chimeras were removed by consensus. Taxonomy was assigned using the SILVA v138.2 [50] and GTDB r220 [51, 52] reference databases and a minimal bootstrap confidence of 50, resulting in a total of 6,342 amplicon sequence variants (ASVs) and 48,516-90,556 read counts per sample (Supplementary Table 3). The DADA2 output was imported without filtering into MicrobiomeAnalyst 2.0 [53] (update 2024-11-01). Data were normalized using total sum scaling. Alpha diversity was calculated at ASV level based on the Shannon index and tested for significance by Welch’s t-test. Beta diversity was calculated at ASV level by principal coordinate analysis (PCoA) based on the Bray-Curtis index and tested for significance by the permutational analysis of variance (PERMANOVA).

For metagenome sequencing, the extracted DNA was sequenced after library preparation with an insert size of 300–400 bp on the DNBSEQ-G400 platform in the paired-end mode with a read length of 150 bp per read. Raw reads were filtered by BGI using SOAPnuke [54] with the parameters “-n 0.001 - l 20 -q 0.5 --adaMis 3 --minReadLen 150” to remove 1) reads matching 50% or more of the adapter sequence with a maximum of 3 base mismatches, 2) reads with a length of < 150 bp, 3) reads with ≥ 0.1% unknown bases (N), 4) reads with > 50% of the bases having a phred quality value < 20. Filtered reads were assembled and binned using a hybrid metaWRAP approach [55]. In brief, reads were first assembled using metaSPAdes v4.0 [56]. Contigs shorter than 1.5 kb were removed. Unused reads were assembled with MEGAHIT v1.2.9 [57], where contigs smaller than 1 kb were removed. The resulting contigs from both assemblies were merged. Contigs were binned employing CONCOCT v.1.1.0 [58], MaxBin2 v2.2.4 [59], and MetaBAT 2 v2.12.1 [60]. Bins were refined and reassembled via metaWRAP v1.3.2 [55]. Quality was assessed using CheckM v1.0.18 [61] (database version 2015_01_16). Bins with a quality score ≥ 50 (completeness – 5x contamination) were considered as metagenome-assembled genomes (MAGs) [62]. MAGs with > 99% ANI were dereplicated using dRep v3.5.0 [63] with default settings but without initial bin filtering. MAGs were taxonomically classified using GTDB-tk v2.4.0 [64] (database version 220) and annotated using Bakta v1.10.4 [65] (database version 5.1).

### Protein analysis

Proteins were extracted from frozen cell pellets and stored at -80 °C, using a harsh disruption including bead-beating, freeze-thaw, and sodium dodecyl sulfate treatment (SDS) as previously described [66] but with some modifications, followed by precipitation using trichloroacetic acid (TCA) and acetone wash as previously described [67] but with some modifications. For this purpose, approximately 250 mg samples were resuspended in 800 µL lysis buffer (1 M Tris-HCl pH 8, 4% (w/v) SDS, 10 mM dithiothreitol (DTT)). Then, samples were subjected to bead-beating at 6.5 m s⁻¹ for 1 min, 95 °C heat treatment at 500 RPM for 20 min, and freezing in liquid nitrogen subsequently. Then, samples were again subjected to bead-beating for 10 min (in 1-minute cycles with cooling of the device between cycles) and heat treatment for 10 min before freezing in liquid nitrogen followed by heat treatment for 10 min, bead-beating for 10 min (in 1-minute cycles with cooling of the device between cycles), freezing in liquid nitrogen, and heat treatment for 10 min. After cell disruption, cell debris and soil particles were removed by centrifugation at 14,000 x g for 10 min at room temperature. To aggregate the proteins, trichloroacetic acid was added to the supernatant from a 100% (w/v) stock solution to a final concentration of 20% (v/v). Samples were mixed gently and incubated at -20 °C overnight. Afterward, samples were centrifuged at 23,000 x g for 30 min at 4 °C and the supernatant was removed. The protein pellet was washed firstly with 800 µL of 0.07% (v/v) β-mercaptoethanol and 1 mM phenylmethylsulfonyl fluoride in acetone, and secondly, with 800 µL 80% (v/v) acetone in water, each followed by centrifugation at 23,000 x g for 30 min at 4 °C. Residual acetone was removed by vacuum centrifugation. The washed protein pellet was resuspended in 30 µL of 100 mM ammonium bicarbonate (AMBIC) buffer (pH 8), supplemented with 5 µL of 2 ng µL⁻¹ bovine serum albumin as internal standard, and adjusted to 5% (w/v) sodium deoxycholate (SDC) by adding 35 µL of 10% (w/v) SDC in water. Cysteine residues were reduced and alkylated by sequentially adding 400 mM DTT to a final concentration of 12 mM (incubated at 37 °C for 30 min at 400 RPM) and 700 mM 2-iodoacetamide to a final concentration of 40 mM (incubated at 20 °C in the dark for 45 min at 400 RPM). Then, samples were diluted with 100 mM AMBIC buffer (pH 8) to a final SDC concentration of 1% (w/v) before adding 6.3 µL of a 0.1 µg L⁻¹ sequencing-grade trypsin stock solution (Promega, Madison, WI, USA). Samples were incubated overnight at 37 °C and 400 RPM for in-solution digestion. Subsequently, neat formic acid was added to a final concentration of 2.5% (v/v) to stop trypsin digestion, and to precipitate SDC as well as undigested proteins. Precipitates were removed by three consecutive centrifugation steps at 16,000 x g for 10 min. The supernatant was desalted using Pierce C-18 tips (Thermo Scientific, Waltham, MA, USA) following the manufacturer’s instructions. The peptides were dried by vacuum centrifugation and stored at -20 °C until analysis. Peptides were resuspended in 30 µL of 0.1% (v/v) formic acid before subjecting 6 µL to Orbitrap mass spectrometry on a nLC-MS/MS system in data-dependent acquisition (DDA) mode as described previously for in-solution digested samples [68]. Mass spectrometry raw data were analyzed using Proteome Discoverer v2.5. In brief, a database search was performed using Sequest HT with the protein database including the dereplicated MAGs derived from metagenome sequencing and universal contaminants [69]. We allowed mass tolerances of ±3 ppm and ±0.1 Da for precursor and fragment ions, respectively and peptide lengths of 7–30 amino acids with a maximum of two missed trypsin cleavages. Oxidation of methionine and carbamidomethylation of cysteine were set as dynamic and static modifications, respectively. Peptide identifications were refined using INFERYS rescoring [70]. The false discovery rate (FDR) was set to 1% at peptide and protein levels using the Percolator [71] and protein FDR validator node. Label-free quantification was performed via the Minora Feature Detector based on precursor ion intensities. Abundances were normalized based on the total peptide amount. Abundance ratios were calculated based on pairwise peptide ratios between sample groups. Significance was tested using a background-based t-test with Benjamini-Hochberg correction. Proteins were considered as reliably identified target proteins when flagged as non-contaminant Master Proteins with at least two peptides identified in at least two replicates of one of the treatments. Reliably identified target proteins with |log2(abundance ratio)| > 1 and adjusted p-value < 0.05 were considered significantly differentially abundant.

### AREc32 *in vitro* bioassay

The AREc32 assay was conducted as described previously [72], with slight modifications. In brief, the compounds were dissolved in Dulbecco’s Modified Eagle Medium (DMEM) supplemented with 10% fetal bovine serum (FBS), serially diluted using a Hamilton Microlab STAR platform (Hamilton, Bonaduz, Switzerland), and added to a 384-well plate containing 2650 cells per well. The plates were incubated at 37 °C for 24 h. Luciferase production was measured via luminescence, after cell lysis and the addition of luciferin and ATP as substrates. Cell viability was assessed using live-cell analysis with the IncuCyte S3 live cell imaging system (Essen BioScience, Ann Arbor, MI, USA). Cytotoxicity was evaluated based on the inhibitory concentration leading to a 10% decrease in cell viability compared to control cells (IC_10_) and its standard error (SE) was calculated from the linear section of concentration-response curves [73] as follows:

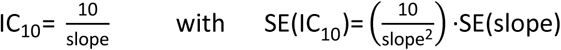

The protocol for the genotoxicity assay (micronucleus formation) can be found in the Supplementary Material.

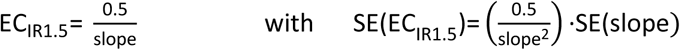

## Results

### Sitagliptin was transformed under nitrate-reducing and methanogenic conditions

To test the potential of microorganisms in hyporheic sediment to transform sitagliptin, we established sediment microcosms using a defined minimal medium with different redox conditions: oxic, nitrate-reducing, sulfate-reducing, and methanogenic. The sediment microcosms were spiked with 700 nM sitagliptin and incubated either aerobically in cotton-plugged Erlenmeyer flasks on a shaker or anoxically in serum bottles with 1 mM KNO_3_, with 1 mM K_2_SO_4_ and 17 mM H_2_, or with only 17 mM H_2_, respectively. Setups with autoclaved sediment and without sediment served as controls for sorption and general abiotic processes. Samples were taken at defined time points and analyzed for sitagliptin, nitrate, sulfate, and methane concentrations. Setups indicating sitagliptin transformation were also analyzed for their microbial community composition.

After incubation for 118–318 days, we observed the decrease of NO_3_^-^ (Supplementary Table 4) and SO_4_^2-^ (Supplementary Table 5) as well as the increase of CH_4_ (Supplementary Table 6) concentrations in the corresponding sediment microcosms. In contrast, the sorption and sediment-free controls showed no relevant reduction of NO_3_^-^ or SO_4_^2-^, nor CH_4_ production, confirming the successful establishment of the targeted conditions. While we did not see significant dissipation of sitagliptin under oxic (Supplementary Table 7) and sulfate-reducing conditions (Supplementary Table 5), sitagliptin dissipated under nitrate-reducing (Figure 1) and methanogenic (Figure 2) conditions.

**Figure 1:**
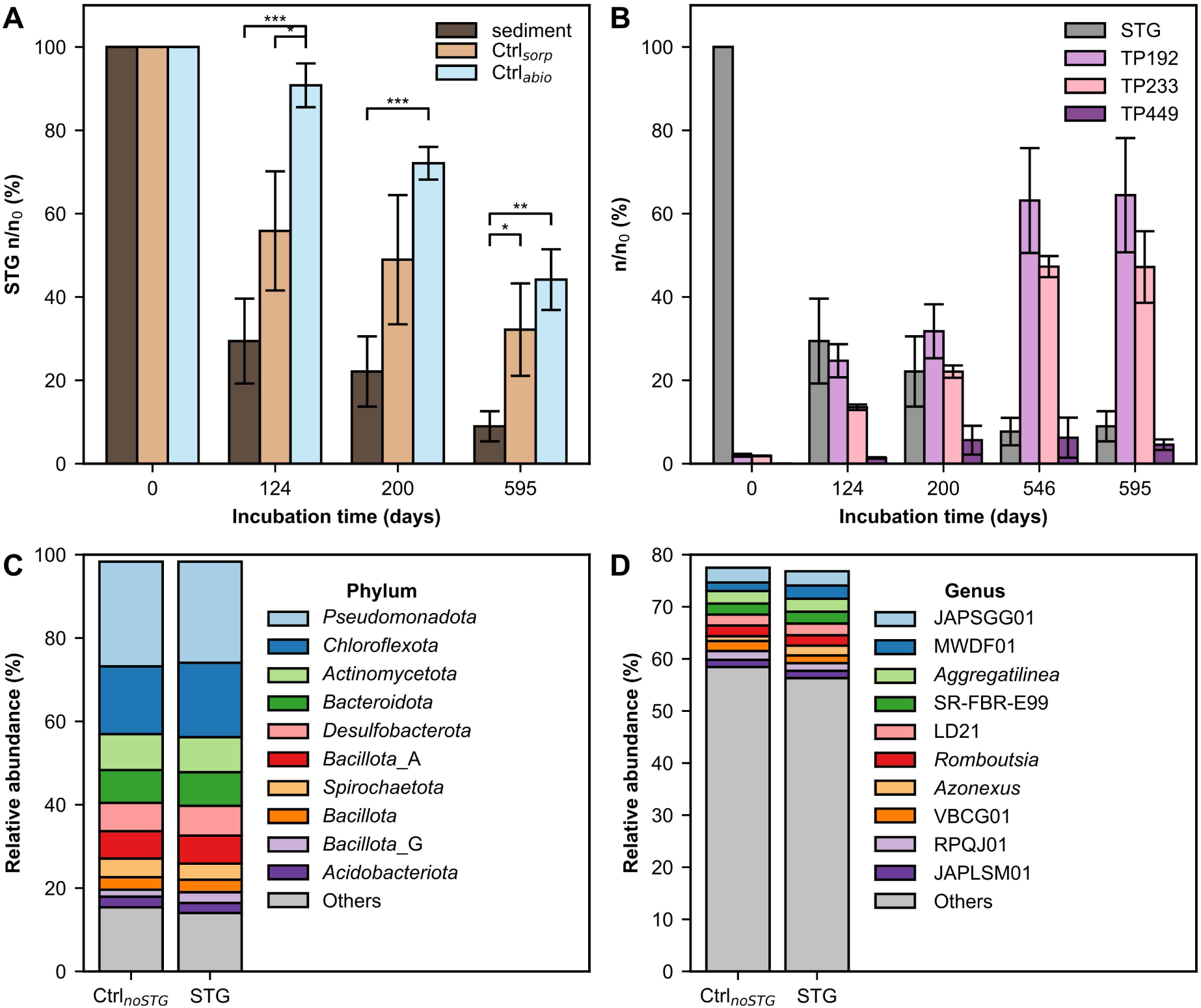
Transformation of sitagliptin (STG) in nitrate-reducing microcosms shown as dissipation of STG compared to control setups with autoclaved sediment (Ctrl_sorp_) or no sediment (Ctrl_abio_) (A) and formation of transformation products in the sediment setup (B). Microcosms were spiked with 700 nM sitagliptin and 1 mM KNO3 at day 0. Relative abundances were calculated in relation to the molar sitagliptin concentration at day 0. The microbial communities in the sediment setups were compared to setups without spike-in after 546 days of incubation based on 16S rRNA gene amplicon sequencing of the v3–v4 region. The 10 most abundant taxa are plotted at phylum (C) and genus (D) level. Taxonomically unclassified amplicon sequence variants were excluded. Values are means of biological triplicates. Error bars denote standard deviations. Statistical differences between setups are indicated with ‘*’, ‘**’, ‘***’ based on an independent Student’s t-test with p < 0.05, p < 0.005, and p < 0.001, respectively.

**Figure 2:**
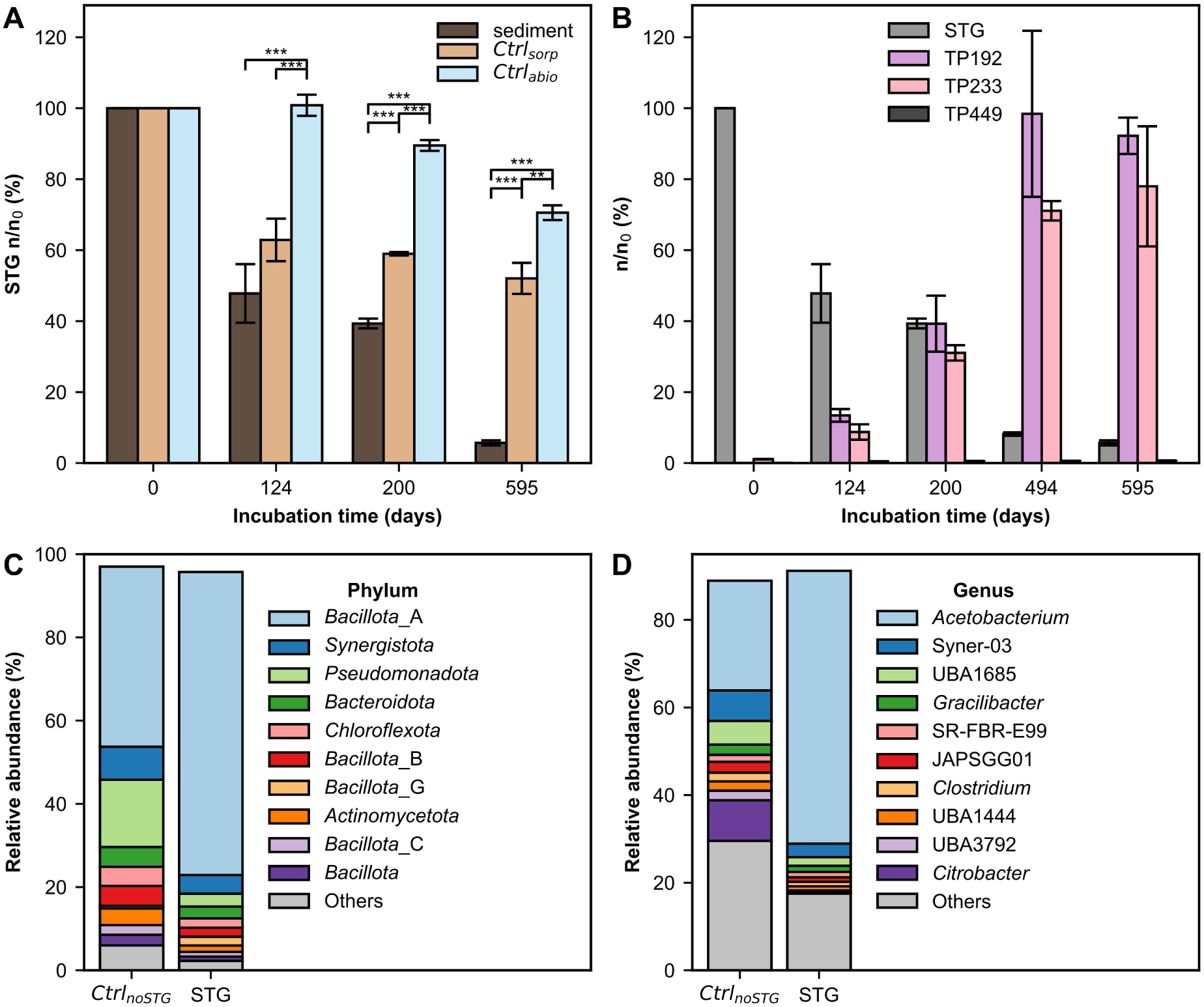
Transformation of sitagliptin (STG) in methanogenic microcosms shown as dissipation of STG compared to control setups with autoclaved sediment (Ctrl_sorp_) or no sediment (Ctrl_abio_) (A) and formation of transformation products in the sediment setup (B). Microcosms were spiked with 700 nM sitagliptin and pressurized with 17 mM H2 at day zero. Relative abundances were calculated in relation to the molar sitagliptin concentration at day 0. The microbial communities in the sediment setups were compared to setups without spike-in after 494 days of incubation based on 16S rRNA gene amplicon sequencing of the v3–v4 region. The 10 most abundant taxa are plotted at phylum (C) and genus (D) level. Taxonomically unclassified amplicon sequence variants were excluded. Values are means of biological triplicates. Error bars denote standard deviations. Statistical differences between setups are indicated with ‘*’, ‘**’, ‘***’ based on Student’s t-test for the means of two independent samples with p < 0.05, p < 0.005, and p < 0.001, respectively.

Most sitagliptin dissipated after 124 days in microcosms with Erpe river sediment incubated under nitrate-reducing conditions (Figure 1A) without a significant increase in cell number (Supplementary Table 8). At all analyzed time points, the sitagliptin concentration in biotic sediment setups was significantly lower than in the sediment-free abiotic control. The concentration of sitagliptin was lower in biotic sediment setups than in sterile sediment setups at all time points, however, the difference between the two setups was only significant after 595 days. In sterile sediment setups, about 50% of sitagliptin was dissipated after 124 days due to sorption and abiotic transformation. In sediment-free setups, about 50% of sitagliptin was dissipated after 595 days due to abiotic processes. We measured the pH after 595 days of incubation: the mean values in sediment microcosms (pH 7.9) were similar to those in the controls (pH 7.6−7.7). Suspect screening revealed the formation of 3 TPs (Figure 1B): TP449, TP233, and TP192, where the number represents the monoisotopic mass of the uncharged TP in Da. For comparison, STG has a monoisotopic mass of 407 Da. TP449 is the N-acetylated form of sitagliptin and was detected at low abundance. TP233 and TP192 result from hydrolysis of the amide bond within sitagliptin and were the major TPs detected under nitrate-reducing conditions. While TP449 was not detected in abiotic controls, TP233 and TP192 were also detected in both abiotic controls, however, to a significantly lower extent (Supplementary Figure 1). To investigate the microbial community compositions, we performed 16S rRNA gene amplicon sequencing of the sediment setups and compared them to controls without sitagliptin spike-in. The majority of the identified organisms belonged to the phyla *Pseudomonadota*, *Chloroflexota*, and *Actinomycetota* (Figure 1C). JAPSGG01, MWDF01, and *Aggregatilinea* were the three most abundantly identified genera (Figure 1D). Notably, the genus *Azonexus* and the families of VBCG01 (*Casimicrobiaceae*) and RPQJ01 (*Steroidobacteraceae*) are known to contain nitrate-reducers. The communities with and without spike-in of sitagliptin did not differ significantly in alpha (Shannon, Welch t-test, p > 0.65) and beta diversity (PCoA, Bray-Curtis, PERMANOVA, p > 0.6).

To assess whether the transformation activity observed was biologically driven and sustainable at higher concentrations, fresh medium containing 50 µM sitagliptin was inoculated with 10% (v/v) of the active microcosms. In the transferred cultures under nitrate-reducing conditions, sitagliptin transformation did not significantly differ from the controls, suggesting that the initial biological activity could not be maintained, and the remaining activity was likely abiotic (Supplementary Figure 1). However, when experiments were conducted with 700 nM sitagliptin and fresh sediment, biotransformation was observed repeatedly in two independent replicate experiments conducted in different years. This indicates that while the activity was not sustainable at high concentrations through transfers, the sediment microbial community consistently demonstrated biotransformation capability at concentrations closer to environmental levels.

Cell numbers significantly increased within 124 days of incubation in microcosms under methanogenic conditions (Supplementary Table 8). About half of the sitagliptin concentration dissipated after 124 days of incubation (Figure 2A). We again observed sorption effects. However, there was significantly (p < 0.001) more sitagliptin dissipation in biotic sediment microcosms compared to sorption controls after 200 days of incubation. We measured the pH after 595 days of incubation: the mean values in biotic sediment microcosms (pH 9.2) differed significantly (p < 0.001) from the controls (pH 7.3-7.4). The amount of acetate and methane increased over time in the biotic sediment microcosms, with methane being constantly more abundant (Supplementary Table 6). Stoichiometry suggests complete conversion of the NaHCO_3_ buffer to acetate and methane after 494 days of incubation, which aligns with the pH increase observed in biotic sediment setups. We mainly detected TP192 and TP233 in biotic sediment setups, closing the mass balance at time points 494 and 595 days (Figure 2B). TP449 was only detected at very low concentrations.

The microbial communities were dominated by the phylum *Bacillota* and its genus *Acetobacterium* (Figure 2C and D). Sitagliptin spike-in led to an increase of *Acetobacterium* (*Bacillota*_A) and a decrease of *Pseudomonadota*, however, the communities did not significantly differ in alpha (Shannon, Welch t-test, p > 0.1) and beta diversity (PCoA, Bray-Curtis, PERMANOVA, p = 0.1). Notably, phyla *Methanobacteriota* and *Halobacteriota* (genera *Methanobacterium*, *Methanothrix*, *Methanoculleus*, and *Methanosarcina*) were the only detected (archaeal) methanogens and had a low relative abundance of 0.01−0.12%.

To assess whether the transformation activity observed under methanogenic conditions was biologically driven and sustainable at higher sitagliptin concentrations, fresh medium containing 50 µM sitagliptin was inoculated with 10% (v/v) of the active microcosms. In the transferred cultures under methanogenic conditions, sitagliptin dissipation remained significantly higher in sediment setups compared to controls, indicating that the biologically induced transformation activity was sustained (Figure 3).

**Figure 3:**
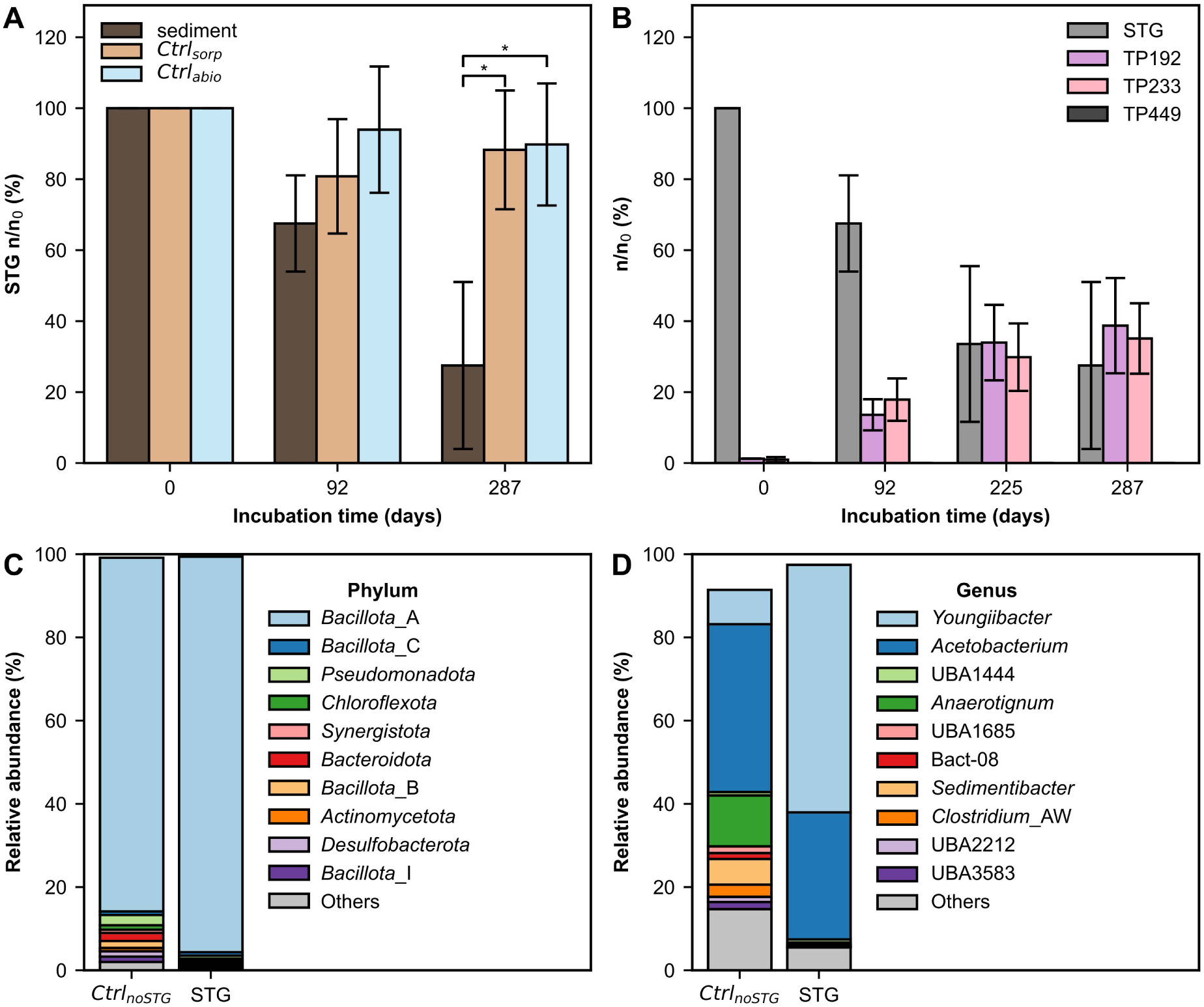
Transformation of sitagliptin (STG) in passed cultures of methanogenic microcosms shown as dissipation of STG compared to control setups with passage from autoclaved sediment (Ctrl_sorp_) or no sediment (Ctrl_abio_) controls (A) and formation of transformation products in the sediment setup (B). Microcosms were spiked with 50 µM sitagliptin and pressurized with 17 mM H2 at day 0. Relative abundances were calculated in relation to the molar sitagliptin concentration at day 0. The microbial communities in the sediment setups were compared to setups without spike-in after 287 days of incubation based on 16S rRNA gene amplicon sequencing of the v3–v4 region. The 10 most abundant taxa are plotted at phylum (C) and genus (D) level. Taxonomically unclassified amplicon sequence variants were excluded. Values are means of biological triplicates. Error bars denote standard deviations. Statistical differences between setups are indicated with ‘*’ based on an independent Student’s t-test with p < 0.05.

The dissipation of sitagliptin after 287 days of incubation was significantly higher in the biotic sediment setups compared to the controls (p < 0.05, Figure 3A). After 378 days of incubation, we measured the pH in the microcosm setups: The mean values in sediment microcosms (pH 9.2) differed significantly (p < 0.05) from the controls (pH 7.3–7.4). While TP192 and TP233 were formed again, TP449 was not detected (Figure 3B). The transfer of the culture to fresh medium led to further enrichment of the phylum *Bacillota* making up more than 90% of the relative abundance (Figure 3C). We detected higher abundance of *Youngiibacter* and lower abundance of *Anaerotignum* and *Sedimentibacter* in microcosms with sitagliptin spike-in compared to controls without sitagliptin (Figure 3D), leading to a significant reduction in alpha diversity (Shannon, Welch t-test, p-value < 0.05).

TP233 and TP192 were previously described as abiotic transformation products of sitagliptin under high pH conditions [74, 75]. To assess whether the observed transformations were dependent on pH, we set up abiotic microcosms with the same medium as the methanogenic setups but without NaHCO_3_. Setups were spiked with 50 µM sitagliptin and the pH was adjusted by the addition of 1M NaOH to pH 7, 8, 9, or 10. The results after 90 days of incubation confirm that sitagliptin undergoes abiotic transformation at pH > 7 (Figure 4). TP192 and TP233 were the only identified TPs. At pH 9, about 40% of the initial spike-in was transformed after 90 days of incubation, which is similar to the observed transformation rate in biotic sediment microcosms incubated under methanogenic conditions.

**Figure 4:**
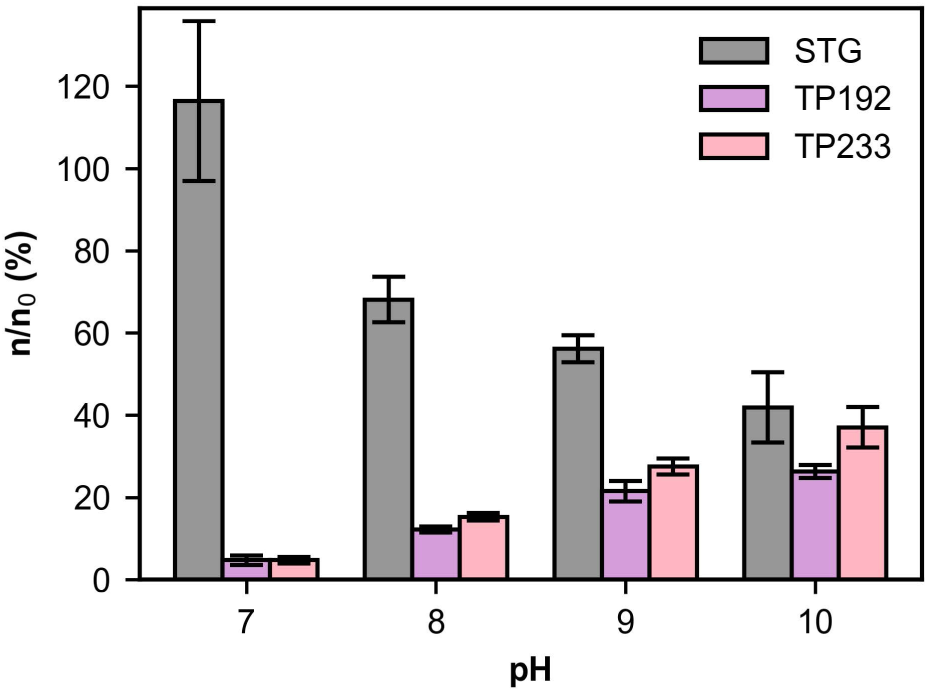
Abiotic transformation of sitagliptin (STG) is dependent on the pH. Concentration of sitagliptin (STG) and its TPs after incubation for 90 days at different pH values. Samples were spiked with 50 µM sitagliptin. Relative abundances were calculated in relation to the molar sitagliptin concentration at day 0. Values are means of biological triplicates. Error bars denote standard deviations.

### TP233 was further transformed under oxic conditions

Under the tested anoxic conditions, no further transformation of TP192 or TP233 was observed. In the hyporheic zone, TrOC and its TPs may be sequentially exposed to different redox zones [76]. Thus, we tested whether the major TPs observed under anoxic conditions are further transformed under oxic conditions. For this purpose, we spiked fresh oxic sediment microcosms with 10 nM, 500 nM, and 10 µM TP192 and TP233, and analyzed samples at designated timepoints for compound dissipation. We did not observe any dissipation of TP192, but a significant (p < 0.001) transformation of 10 nM TP233 with a removal of 92% within 7 days (Figure 5).

**Figure 5:**
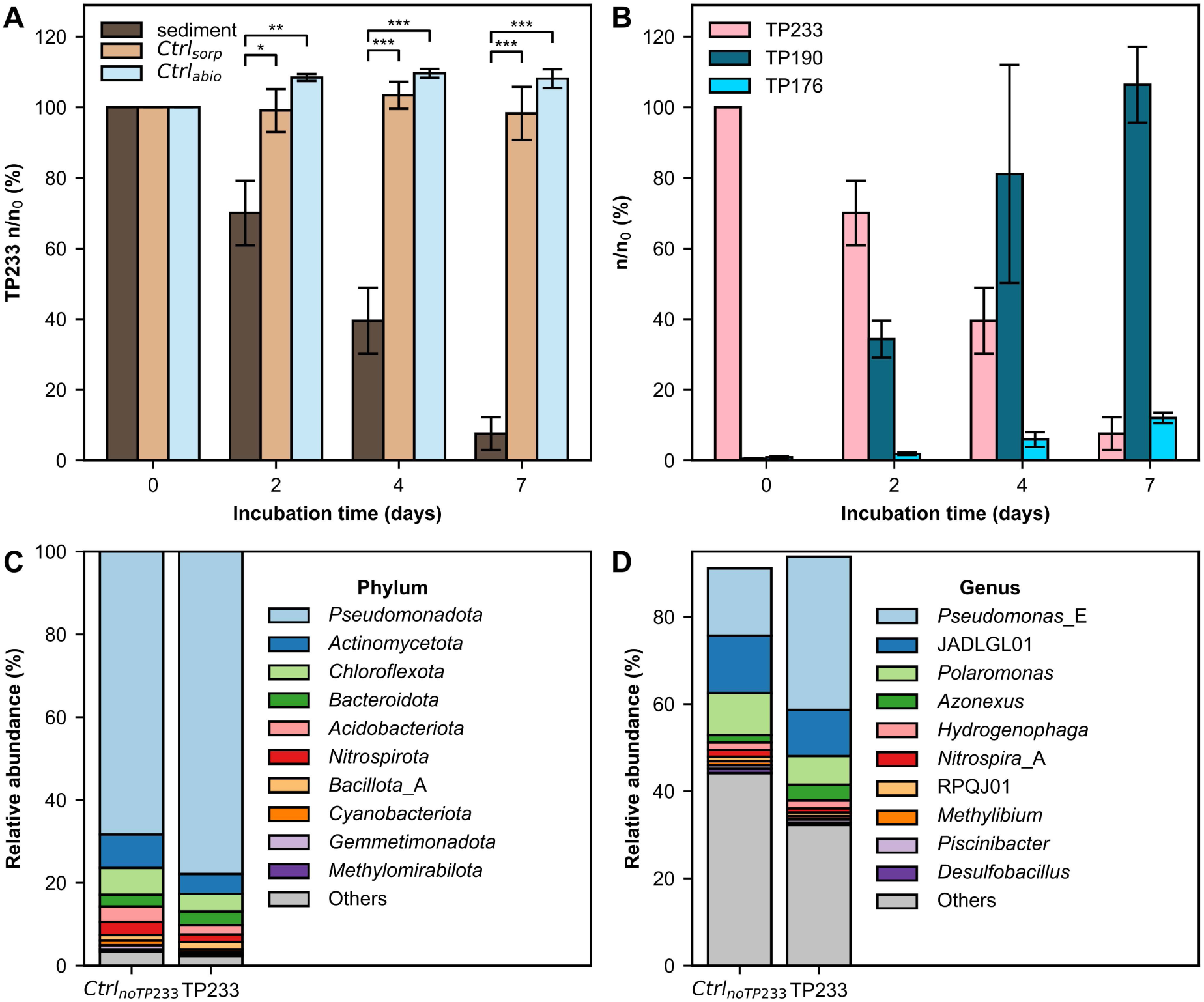
Transformation of TP233 in oxic sediment microcosms shown as dissipation of TP233 compared to autoclaved sediment (Ctrl_sorp_) or no sediment (Ctrl_abio_) controls (A) and formation of TP190 and TP176 in the sediment setup (B). Microcosms were spiked with 10 nM of TP233 at day 0. Relative abundances were calculated in relation to the molar TP233 concentration at day 0. The microbial communities in the sediment setups were compared to setups without spike-in after 7 days of incubation based on 16S rRNA gene amplicon sequencing of the v3–v4 region. The 10 most abundant taxa are plotted at phylum (C) and genus (D) level. Values are means of biological triplicates. Taxonomically unclassified amplicon sequence variants were excluded. Error bars denote standard deviations. Statistical differences between setups are indicated with ‘*’, ‘**’, ‘***’ based on Student’s t-test for the means of two independent samples with p < 0.05, p < 0.005, and p < 0.001, respectively.

Suspect screening revealed the sequential formation of the major TP190 (2-(2,4,5-trifluorophenyl)acetic acid) and the minor TP176 (2,4,5-trifluorobenzoic acid), both confirmed using authentic reference standards (Figure 5B). We did not detect any transformation of TP233 in sorption and abiotic controls, indicating biotransformation of TP233. TP190 could have been formed by oxidative deamination of TP233 with subsequent oxidation and cleavage of acetate. TP176 could have been formed by the subsequent oxidation of TP190 with subsequent cleavage of formate.

The microbial communities were dominated by the phylum *Pseudomonadota* (Figure 5C) and the genera *Pseudomonas*, JADLGL01 (classified as *Labrys* by SILVA v138.2), and *Polaromonas* (Figure 5D). We did not detect significant differences in alpha (Shannon, Welch’s t-test, p > 0.28) and beta diversity (PCoA, Bray-Curtis, PERMANOVA, p = 0.3) between samples with and without TP233 spike-in.

To further explore the functional capabilities and activity of the identified microorganisms, we performed metagenome-resolved metaproteomics. For this purpose, we sequenced DNA extracts from the microcosms after 7 days of incubation as well as from frozen sediment that had been used as the inoculum for the microcosms. Assembly, binning, and dereplication at 99% average nucleotide identity yielded 43 unique metagenome-assembled genomes (MAGs) from an initial set of 78 MAGs (Supplementary Table 10). The unique MAGs corresponded to the highly abundant organisms previously identified through 16S rRNA gene amplicon sequencing. Metagenome-resolved metaproteomics reliably identified 1,692 target proteins (Supplementary Table 11) of the 43 MAGs, confirming their existence and activity. JADLGL01, *Pseudomonas* (*asiatica*), and *Polaromonas* emerged as the most abundant genera at protein level — consistent with the 16S rRNA gene amplicon sequencing results (Supplementary Table 12).

We then tested the identified proteins for differential abundance between microcosms spiked with 10 nM TP233 and controls. Of the 1,692 identified proteins, 39 were exclusive to TP233 samples, and 8 were exclusive to controls. Additionally, 100 proteins were significantly more abundant, while 22 were significantly less abundant in the TP233 samples. Notably, 110 of the 139 (79%) proteins identified as exclusive or significantly more abundant in TP233 samples were attributed to MAGs classified as *Pseudomonas asiatica* (Supplementary Table 11). Among the 110 significantly more abundant *P. asiatica* proteins, several were related to oxidative deamination, stress response, and tricarboxylic acid (TCA) cycle. Specifically, the monoamine oxidase of *P. asiatica* was exclusively identified in TP233 samples. Monoamine oxidases catalyze the oxidative deamination of amino groups to keto groups, releasing H_2_O_2_ and NH_3_. Likewise, H_2_O_2_ and NH_3_-related stress proteins of *P. asiatica*, including alkylhydroperoxidase, catalase/peroxidase, thiol peroxidase, glutamate—ammonia ligase, and universal stress proteins were significantly more abundant in TP233 samples than in controls. In addition, several *P. asiatica* proteins related to the tricarboxylic acid (TCA) cycle, including dihydrolipoyl dehydrogenase, succinate dehydrogenase, 2-oxoglutarate dehydrogenase, and pyruvate dehydrogenase, were significantly more abundant in TP233-treated samples compared to controls. Notably, formate dehydrogenase H from *Methylibium petroleiphilum* was exclusively identified in TP233 samples.

### Sitagliptin transformation in distinct microbial communities shaped by redox conditions

To compare the microbial communities enriched under distinct redox conditions, we performed a principal coordinate analysis of the beta diversity based on the Bray-Curtis index of the different microcosms and assessed the corresponding alpha diversity based on the Shannon index (Figure 6).

**Figure 6:**
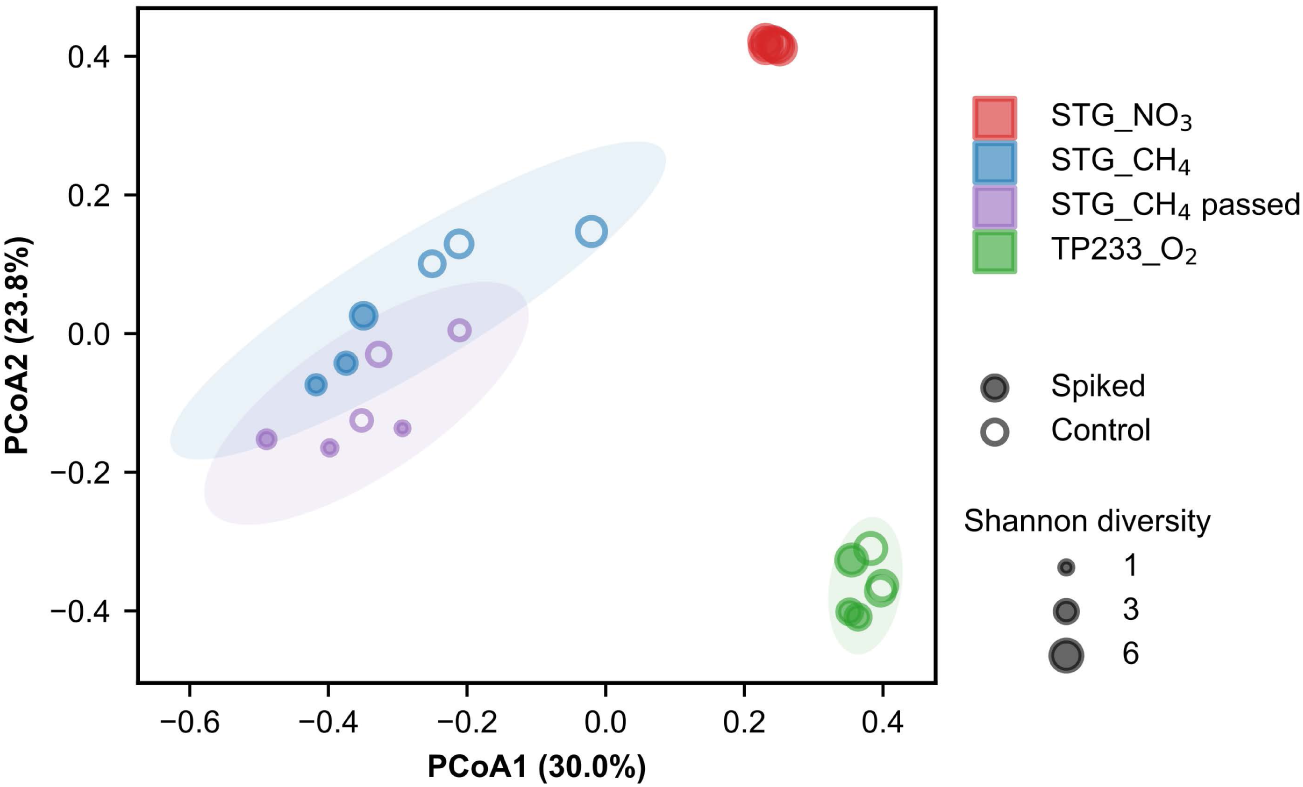
Principal coordinate analysis (PCoA) showing differences in beta diversity based on the Bray-Curtis index between microcosms transforming sitagliptin (STG) under nitrate (NO_3_ ^-^) reducing and methanogenic (CH_4_) conditions and TP233 under oxic (O_2_) conditions. Alpha diversity based on the Shannon index is depicted as dot area. Community compositions were assessed by 16S rRNA gene amplicon sequencing of the v3–v4 region. Diversity measures were calculated at the level of amplicon sequence variants. Ellipses indicate 95% confidence intervals assuming a multivariate normal distribution.

Communities detected under different redox conditions exhibited significant differences in both alpha diversity (ANOVA, p < 0.001) and beta diversity (PERMANOVA, p = 0.001). Alpha diversity and similarity between samples were highest in NO₃⁻-reducing conditions and lowest in methanogenic conditions. The impact of redox conditions on beta diversity exceeded that of sitagliptin or TP233 spike-in.

### Sitagliptin and its TPs were detected in the River Erpe

To test whether the transformation processes observed in batch cultures are relevant in the hyporheic zone, we analyzed porewater samples from the River Erpe. Specifically, we took samples across a depth profile at two distinct sampling sites characterized by contrasting sediment compositions. Site A consisted predominantly of muddy sediment, while Site B was characterized by sandy sediment. At each site, four independent porewater samplers were deployed in close proximity to account for local heterogeneity. Samples were analyzed by LC-MS/MS via suspect screening (Figure 7).

**Figure 7:**
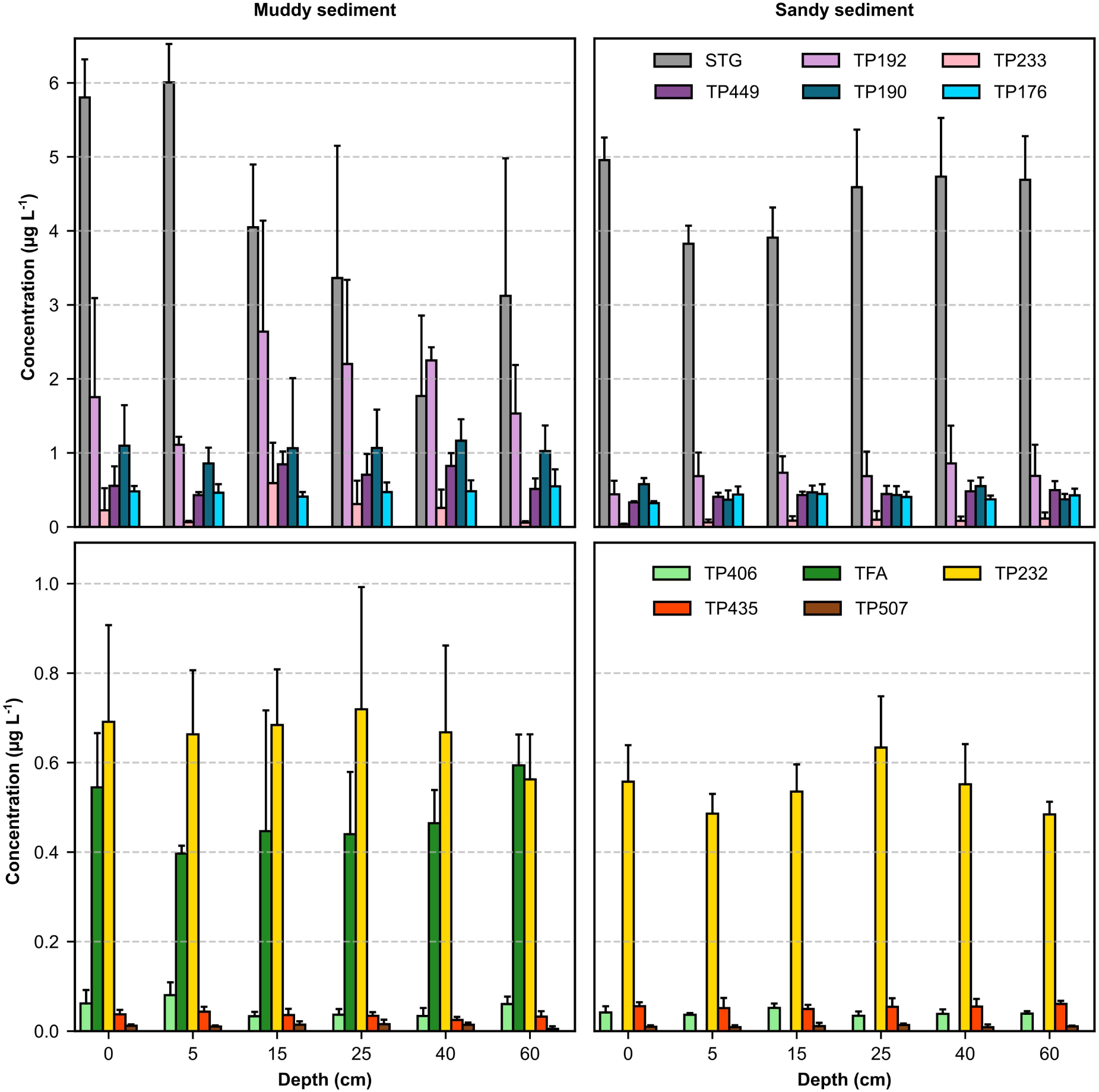
Occurrence of sitagliptin (STG) and its transformation products (TPs) in porewater of the River Erpe across a depth profile at two different sites. Site A was characterized by muddy sediment composition while Site B exhibited predominantly sandy sediment characteristics. Values represent means + standard deviation from replicate samplers (n=4) deployed within each site. Substances in the upper subplots were also identified in batch experiments. Substances in the lower subplots were exclusively detected in the River Erpe. Note the differences in the scale of the y-axes. Trifluoroacetic acid (TFA) was below the limit of quantification (0.05 µg L⁻¹) in samples from Site B.

At both sampling sites and across all sediment depths, we identified sitagliptin and all TPs observed in previous batch experiments (Figure 7A and B). Additionally, we detected and quantified TP406 (1-[3-(trifluoromethyl)-6,8-dihydro-5H-[1,2,4]triazolo[4,3-a]pyrazin-7-yl]-4-(2,4,5-trifluorophenyl)butane-1,3-dione), a previously reported sitagliptin TP [11], and trifluoroacetic acid (TFA), a putative environmental end product of sitagliptin transformation, using reference standards. Furthermore, we detected TP232, TP435, and TP507 in positive ionization mode based on characteristic MS/MS transitions. TP232 was detected with an *m/z* of 233.0 and the characteristic fragments with *m/z* ratios of 175.0, 155.0, and 127.0. TP435 was detected with an *m/z* of 436.1, and characteristic fragments with a *m/z* of 244.1 and 193.1. TP507 was detected with an *m/z* of 508.1 and characteristic fragments with *m/z* ratios 391.1, 193.1, and 174.1. Concentrations of TPs without reference standards were estimated semiquantitatively using calibrations of structurally similar TPs sharing a fragment ion.

Sitagliptin concentrations did not significantly differ between the muddy and the sandy site when combining the concentrations measured across all sampling depths. However, TP192, TP233, TP449, TP190, TP176, TFA, and TP232 concentrations were significantly higher (Welch’s t-test with Benjamini-Hochberg correction, p < 0.05) at the muddy site compared to the sandy site, while TP435 concentration was significantly lower (p < 0.001). Depth had no significant effect on TP concentrations (one-way ANOVA with Benjamini-Hochberg correction, p > 0.11). However, we observed a site-specific effect of depth on sitagliptin concentrations: In porewater from the muddy site, sitagliptin concentrations varied significantly with depth (p < 0.01), peaking at 6.0 ± 0.5 µg L⁻¹ at 5 cm and decreasing to 1.8 ± 0.9 µg L⁻¹ at 40 cm. At the sandy site, STG concentrations did not differ significantly with depth (p > 0.25).

We analyzed the correlation between the measured sitagliptin and TP concentrations with ion concentrations characteristic for anoxic conditions from the same sampling campaign (Gerundt et al., unpublished manuscript, 2025). In addition to the electron acceptors NO_3_ ^-^ and SO_4_ ^2-^ tested in this study, we also included Fe^2+^, indicative for anaerobic Fe^3+^ reduction in the hyporheic zone [77]. At the muddy site, we identified significant correlations of STG and its TPs with NO_3_ ^-^, SO_4_ ^2-^, and Fe^2+^ (Figure 8).

**Figure 8:**
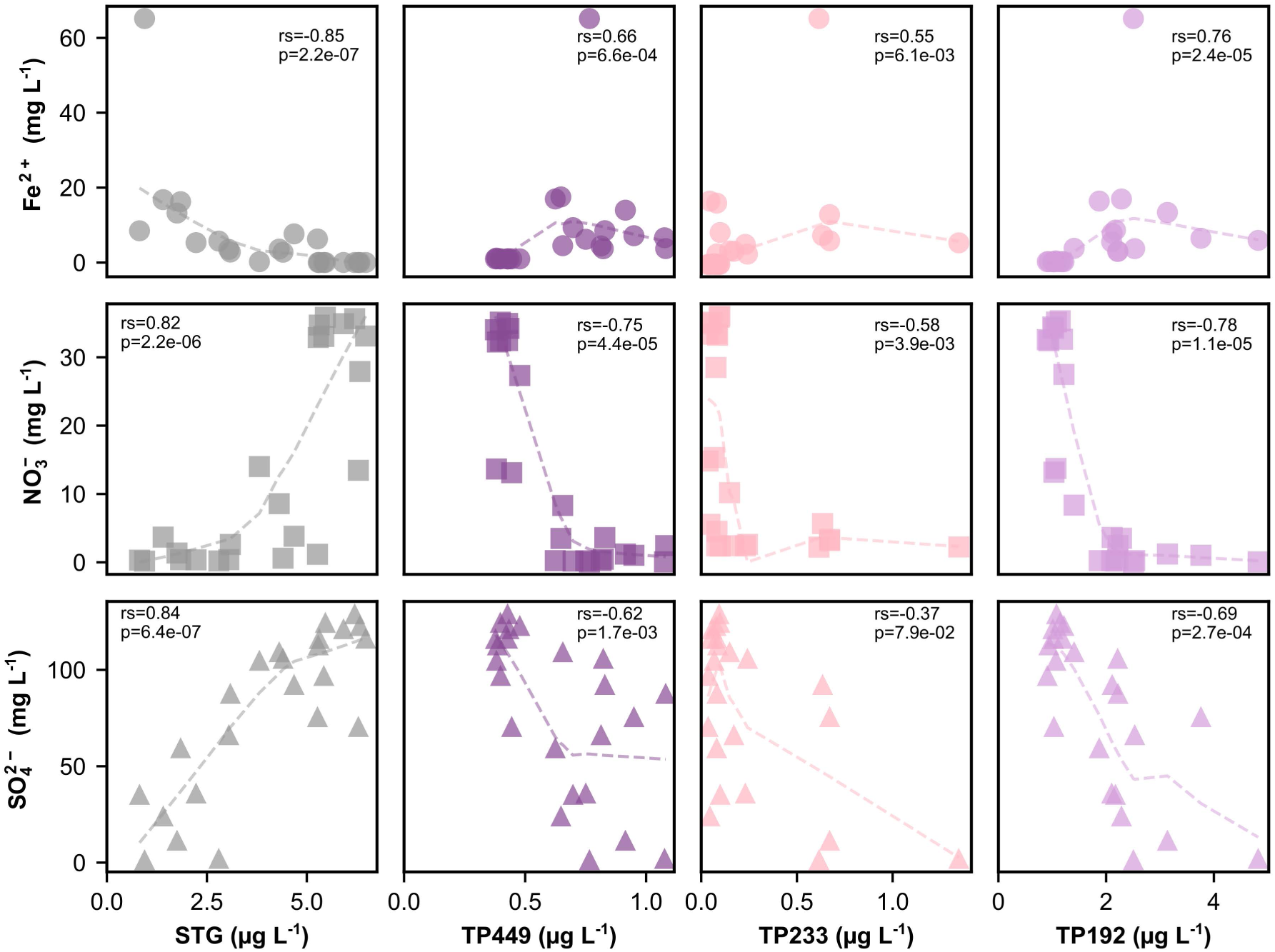
Correlation of sitagliptin (STG), TP449, TP233, and TP192 with iron(II) (Fe²⁺), nitrate (NO₃⁻), and sulfate (SO₄²⁻) concentrations in pore water samples collected from the River Erpe at Site A (muddy). Dashed lines represent trend lines based on locally weighted scatterplot smoothing (LOWESS) with a smoothing factor of 0.7. Spearman’s rank correlation coefficients (rs) and corresponding p-values are shown in each subplot.

Sitagliptin exhibited positive correlations with NO₃⁻ (both sites) and SO₄²⁻ (muddy site only), while negatively correlating with Fe²⁺ at both sites. Conversely, TP449, TP233, and TP192 showed negative correlations with NO₃⁻ and positive correlations with Fe²⁺ at both sites, as well as with SO₄²⁻ at the muddy site. In general, correlations were stronger at the muddy site compared to the sandy one (Supplementary Figure 2).

### Sitagliptin TPs were less toxic than the parent compound

To assess the toxicity of sitagliptin and its TPs, we performed the AREc32 *in vitro* bioassay, a reporter gene assay for cytotoxicity and oxidative stress response (Figure 9) [72].

**Figure 9:**
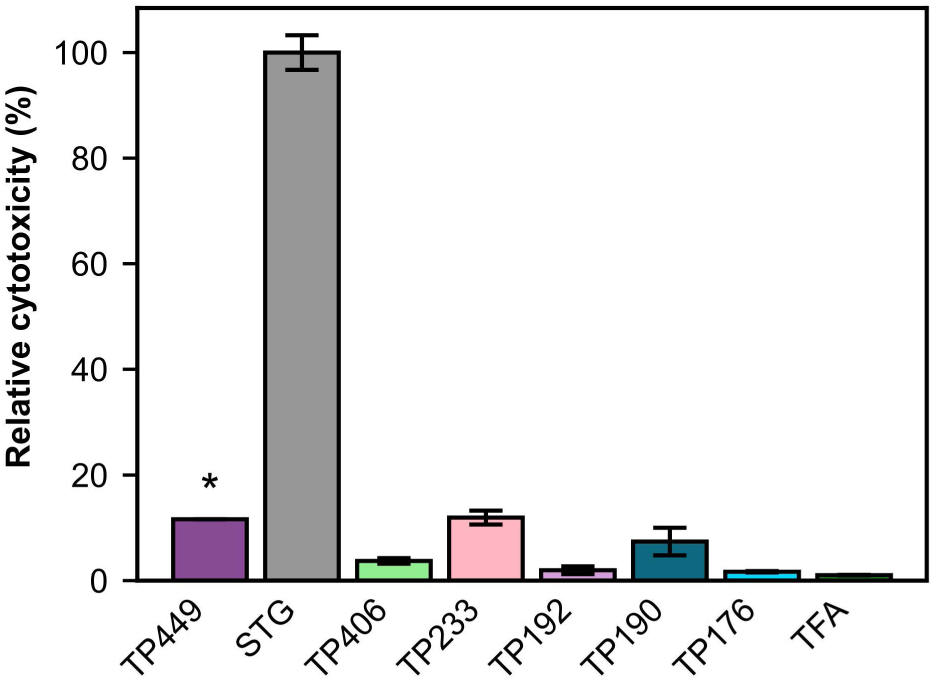
Relative cytotoxicity of sitagliptin (STG), its transformation products (TPs), and trifluoroacetic acid (TFA) in the AREc32 *in vitro* bioassay expressed as relative effect potency REP = IC10(STG)/IC10(TP). Error bars denote standard errors. ‘*’ indicates that REP < 12% for TP449 as the highest tested concentration did not reach 10% cytotoxicity.

All commercially available TPs exhibited lower cytotoxicity than sitagliptin. Furthermore, TP190 and TP176 showed lower cytotoxicity than their parent compound TP233. Only sitagliptin and TP192 induced an oxidative stress response, however, the effect was of low specificity and associated with cytotoxicity (Supplementary Figure 3B). We also evaluated cytotoxicity and genotoxicity via the *in vitro* mammalian cell micronucleus test (Supplementary Table 9). Cytotoxicity results from the micronucleus test were consistent with those from the AREc32 bioassay (Supplementary Figure 3A). Only TP406, TP233, and TP176 induced micronucleus formation, however, the observed effect was non-specific and occurred at concentrations close to 50% cytotoxicity (Supplementary Figure 3C).

## Discussion

Our results indicate that sitagliptin undergoes sequential and redox-dependent transformations in urban hyporheic zones leading to different transformation products (Figure 9).

**Figure 9:**
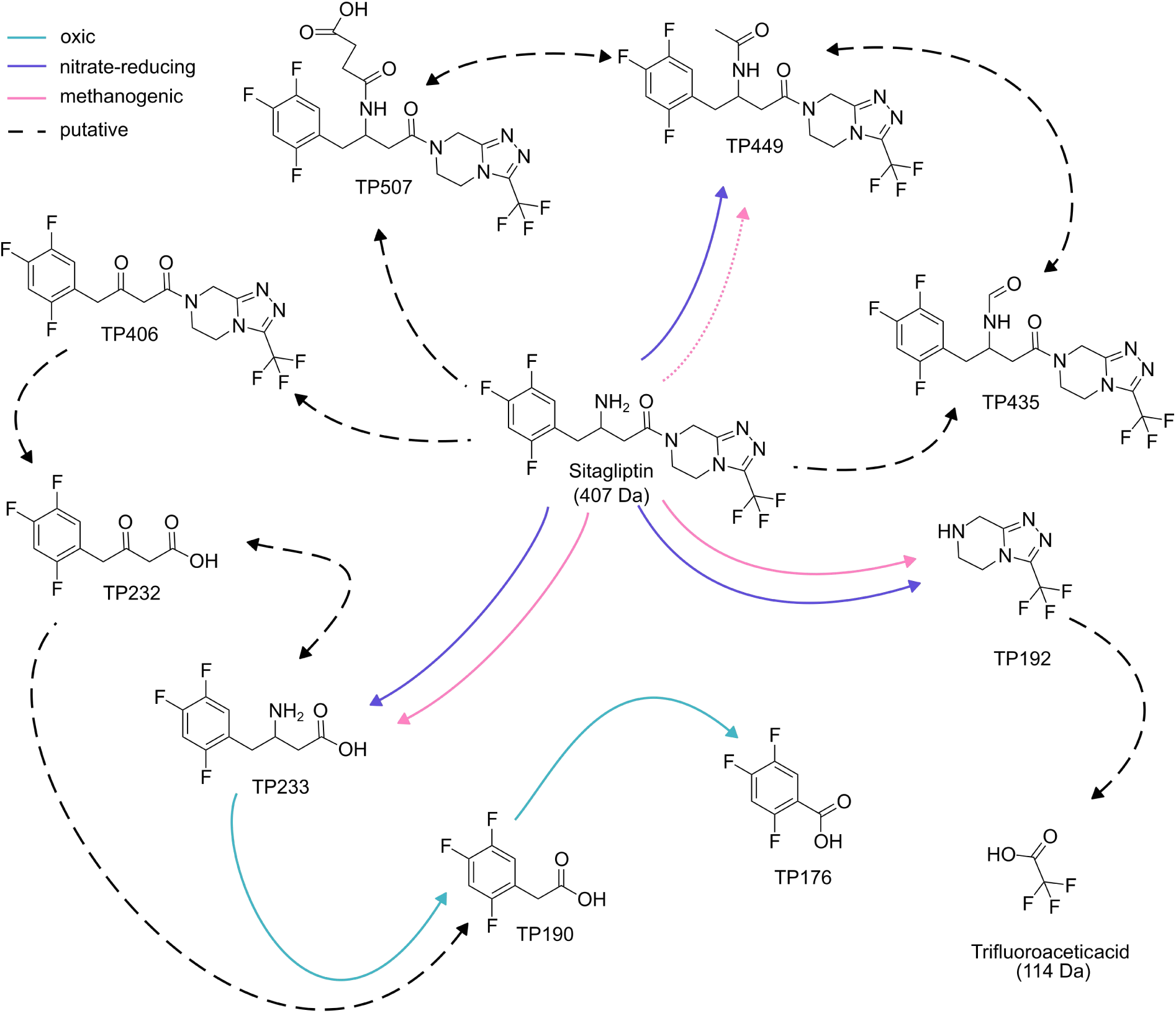
Overview of observed sitagliptin transformation products in the hyporheic zone of the River Erpe, Germany. The reaction pathways are colored according to the redox conditions of the batch microcosms in which the transformation products were observed. Dotted lines indicate a very low amount of transformation. Dashed lines are putative reaction pathways based on TP identifications in pore water samples of the River Erpe but not observed in batch microcosms.

### Attenuation of sitagliptin under different redox conditions

Sitagliptin was hydrolyzed at its amide bond in microcosms under methanogenic and nitrate-reducing conditions, with significantly greater transformation observed in biotic setups compared to abiotic controls, indicating microbial involvement in both environments. Under methanogenic conditions, sitagliptin hydrolysis likely occurred indirectly, catalyzed by a microbially induced pH increase, as suggested by pH-dependent abiotic hydrolysis observed in our experiments and previous studies [74, 75]. Conversely, under nitrate-reducing conditions, no significant pH elevation was detected, suggesting a distinct, pH-independent transformation pathway. A pH-independent pathway could be attributed to co-metabolic transformation, however, co-metabolic hydrolysis of tertiary amides (such as sitagliptin) typically requires initial N-dealkylation [78], and we did not detect any N-dealkylation products (opening of the piperazine ring), consistent with prior research [11]. Direct amide hydrolysis of tertiary amines could also be catalyzed by microbial enzymes like the DEET hydrolase (*dthA*) [79, 80]. Previous studies also demonstrated abiotic hydrolysis of sitagliptin’s amide bond under strongly reducing conditions using at least 10 mM titanium(III) citrate [81]. In contrast, our experiments under both nitrate-reducing (no L-cysteine added) and methanogenic conditions (4 mM L-cysteine) showed no substantial abiotic transformation. The weaker reducing potential of L-cysteine (E°’= -0.21 V [82]) compared to titanium(III) citrate (E°’ = -0.48 V [83]) suggests that more negative redox conditions are required for abiotic reductive amide hydrolysis of sitagliptin.

Sitagliptin hydrolysis generated TP233 and TP192, while subsequent oxic conditions facilitate TP233’s further degradation to TP190 and TP176. This highlights that alternating redox conditions can promote the degradation of TrOCs, as shown previously for sulfamethoxazole [46], halogenated organics [84], dioxanes [85], or azo dyes [86]. Henning et al. (2019) hypothesized the formation of TP232 and TP233 as initial transformation products of sitagliptin that they did not detect [11]. Our study provides experimental evidence for TP233 formation and its role as a precursor for TP190 and TP176 formation. Although we did not detect TP232 directly, our metaproteomic data also support the hypothesis that oxidative deamination of TP233 results in TP232 as an intermediate (see for a detailed discussion below).

A second observed transformation pathway proceeded via N-acetylation of sitagliptin and occurred mainly under nitrate-reducing conditions and to a very low degree under methanogenic conditions, leading to the formation of TP449. Henning et al. (2019) previously identified sitagliptin transformation product TP449 besides TP507, TP435, TP406, TP192, TP190, and TP176 in aerated systems using carrier-attached biomass and activated sludge [11]. The observed formation of similar transformation products in aerated systems and anoxic microcosms might be explained by the presence of anaerobic microsites despite aeration [87, 88] or enzymatic reactions independent of molecular oxygen. Previous studies reported a resistance of sitagliptin to transformation by cytochrome P450 [14], a key enzyme in the aerobic transformation of multiple antidiabetics, further supporting the involvement of processes independent of molecular oxygen in the first steps of sitagliptin transformation.

Sorption behavior of sitagliptin varied between redox conditions, and was primarily observed in nitrate-reducing and methanogenic microcosms. Sorption could be differentiated from transformation by autoclaved controls and the absence of transformation products. We did not observe significant sorption of sitagliptin under oxic and sulfate-reducing conditions. The observed redox-dependent variation in sorption behavior could be attributed to the distinct physicochemical properties of sediments from different depths for each experimental condition. We experimentally determined a p*K*_a_ value of 8.16 for sitagliptin, indicating that it exists partially in the cation form in the environment. Cations typically adsorb more strongly than their neutral counterparts to clay and organic carbon-rich soils [89]. Likewise, it has been shown that sorption of sitagliptin correlates positively with soil organic matter content and negatively with pH [90].

The environmental relevance of our batch experiments is supported by the detection of the same TPs in the River Erpe pore water and the correlation of TP192, TP233, and TP449 with redox-indicative Fe^2+^, NO_3_ ^-^, and SO_4_ ^2-^ ions. The observed amide hydrolysis proceeded slowly in batch experiments under anoxic conditions, with half-lives of approximately 100 days. However, attenuation rates in the field can exceed those in batch experiments by an order of magnitude, suggesting potentially faster transformation in environmental systems [81]. Our field experiments revealed spatial heterogeneity in sitagliptin transformation in hyporheic zones. Specifically, muddy sites showed enhanced transformation of sitagliptin compared to sandy sites. Muddy sites were characterized by higher organic content and more consistent anoxic conditions than sandy sites (Gerundt et al., unpublished manuscript, 2025). Hydrological modeling indicated increased transit times in muddy sites, suggesting that a combination of elevated transit time and distinct redox conditions contributed to the enhanced transformation (Gerundt et al., unpublished manuscript, 2025). Hyporheic fluxes and transit times could be further increased, *e.g.*, by employing surface-subsurface structures in the river bed [91]. Sitagliptin TP concentrations also varied along depth profiles within individual samplers, reflecting the spatial heterogeneity of the hyporheic zone with biogeochemical hot spots [92], preferential flow paths [93, 94], and microsites [95, 96]. We detected TFA, a persistent transformation product of concern [97], at one site in the River Erpe, but not in our microcosm experiments, indicating no biotic transformation of sitagliptin to TFA. While TFA can originate from various sources, it may also form via photolysis from aromatic compounds containing −CF_3_ moieties, such as sitagliptin or TP192 [98]. Since our microcosms were incubated in the dark, further research should investigate the photolytic formation of TFA from sitagliptin and its TPs.

### Sitagliptin-transforming microbiomes

Microbial community analyses of batch microcosms indicated that redox conditions, rather than exposure to sitagliptin, drive community structures, which is in accordance with field observations that redox gradients strongly influence microbial community structures in hyporheic zones [99]. The impact on microbial community composition could be more pronounced at sitagliptin concentrations higher than those used in this study, particularly if metabolic processes are involved. The accumulation of acetate and methane in the methanogenic microcosms suggests complete consumption of the NaHCO_3_ buffer by two microbial processes: methane production by hydrogenotrophic (*Methanobacterium*) and acetoclastic (*Methanothrix*, *Methanoculleus*, and *Methanosarcina*) methanogens, and acetate production by hydrogenotrophic acetogens (*Acetobacterium*). The buffer depletion could have led to the observed pH increase, potentially contributing to sitagliptin hydrolysis [100–102]. Despite methane being more abundant than acetate at all time points, 16S rRNA gene amplicon sequencing showed hydrogenotrophic acetogens exceeding methanogens by at least two orders of magnitude. The apparent discrepancy between metabolite concentrations and microbial abundances could be attributed to the use of universal primers targeting the V3–V4 hypervariable region, often underestimating archaeal abundances [103, 104]. In oxic TP233-degrading microcosms, we predominantly detected the genera *Pseudomonas*, JADLGL01 (classified as *Labrys* by SILVA v138.2), and *Polaromonas*. These genera possess diverse metabolic capabilities and are known for aromatic compound degradation, potentially enabling the transformation of TP233 to TP190 and TP176 [105–107]. *Labrys* and *Pseudomonas* species are reported to degrade organofluorine compounds [106, 107] and to perform PFAS chain shortening [108, 109].

Metaproteomics revealed significantly higher abundances of mainly *P. asiatica* proteins in TP233-transforming cultures, including a monoamine oxidase, stress-response proteins, and proteins involved in the TCA cycle. This suggests that *P. asiatica* is a player in the oxidative deamination of TP233 to TP232, followed by beta-oxidation, releasing TP190 and acetate. Subsequently, acetate could be metabolized by *P. asiatica* via the TCA cycle. TP190 could undergo alpha-oxidation to TP176, releasing formate. Formate may be further oxidized by formate dehydrogenase H from *M. petroleiphilum*, which was exclusively identified in the TP233 samples. The proposed degradation pathway could be further tested via stable isotope probing [110]. However, stable isotope probing would require sitagliptin and TP233 isotopologues, that are currently commercially unavailable and costly to synthesize.

### Toxicology of sitagliptin TPs

All investigated TPs exhibited lower toxicity than sitagliptin on the tested cell lines, with cytotoxicity levels matching previous *in vitro* assays of sitagliptin [35–37] and consistent with prior findings for sitagliptin TPs [38, 39]. Sequential anaerobic/aerobic conditions in the hyporheic zone appeared to facilitate sitagliptin detoxification, corroborating earlier findings where anaerobic pretreatment reduced wastewater toxicity [111]. This suggests that alternating redox conditions promote both trace organic compound degradation and detoxification. Measured sitagliptin concentrations in the River Erpe were three orders of magnitude below reported no-observed-effect concentrations for green algae, *Daphnia magna*, and *Pimephales promelas* in 3−33 day exposure studies, suggesting no acute ecological risk [23]. However, the high concentrations of sitagliptin TPs detected in the River Erpe, combined with typically encountered additive mixture effects, may contribute to the overall toxicity of wastewater treatment plant effluents [112, 113]. Moreover, potential chronic effects from long-term exposure remain unexplored, and several TPs detected in pore water samples lack toxicological characterization due to unavailable reference standards. In addition, slowly degrading compounds and non-degradable transformation products may accumulate in partially closed water cycles, such as in Berlin, Germany, over time and eventually reach (eco)toxicologically relevant levels [15, 16, 114].

Overall, our study highlights the natural biotransformation potential of sequential anaerobic/aerobic conditions in urban hyporheic zones and contributes parameters for engineering hyporheic zones for enhanced contaminant removal in receiving waters. Furthermore, our research emphasizes the importance of including transformation products in risk assessments, particularly given that most oxidative stress response in water samples originates from unknown chemicals [115]. By this, our findings contribute to an overall deeper understanding of TrOC and PFAS transformation and highlight the important role of the hyporheic zone in micropollutant attenuation.

## Availability of data and materials

Raw sequencing reads and dereplicated MAGs with a completeness of at least 90% have been deposited at SRA and GenBank [116], respectively, with the BioProject identifier PRJNA1230214 (reviewer link: https://dataview.ncbi.nlm.nih.gov/object/PRJNA1230214?reviewer=7gcamt6o4ehjav6 <u>u6b6nv8tphh</u>). The mass spectrometric proteomics data have been deposited to the ProteomeXchange Consortium (http://proteomecentral.proteomexchange.org) via the PRIDE partner repository [117] with the dataset identifier PXD061324 and 10.6019/PXD061324 (reviewer token: WKjdU9RvppgK). Dose-response curves from *in vitro* bioassays are available in the Supplementary Material (Supplementary Figure 4 & Supplementary Figure 5). Any other relevant data will be made available upon request

## Acknowledgements

We thank Alexander Kunze, Lasse Kürschner, and Ines Mäusezahl for assistance in cultivation and sample preparation, and Matthias Bernt for maintaining the UFZ’s Galaxy instance. Furthermore, we acknowledge support from Jörg Lewandowski and Christoph Reith in field sampling, Chang Ding in metagenomics, Benjamin Scheer, Jimmy Köpke, and Caglar Akay in LC-MS/MS, Steffen Kümmel in GC, and Anne Knoll and Kerstin Ethner in IC measurements. Metagenome analysis was performed at the High-Performance Computing (HPC) Cluster EVE, a joint effort of both the Helmholtz Centre for Environmental Research – UFZ (http://www.ufz.de/) and the German Centre for Integrative Biodiversity Research (iDiv) Halle-Jena-Leipzig (http://www.idiv-biodiversity.de/). We would like to thank Thomas Schnicke, Ben Langenberg, Guido Schramm, Toni Harzendorf, Tom Strempel, Lisa Schurack, and Christian Krause for maintaining EVE. Metaproteomic data was acquired at UFZ at the Centre for Chemical Microscopy (ProVIS), which is supported by the European Regional Development Fund (EFRE) and the Helmholtz Association. We gratefully acknowledge access to the platform CITEPro (Chemicals in the Environment Profiler) funded by the Helmholtz Association for bioassay measurements and financial support from the Helmholtz POF IV Topic 9 “Healthy Planet-towards a non-toxic environment”. We thank Caroline Gebert, Sophia Bardehly, Jenny Braasch, Julia Huchthausen, Niklas Wojtysiak, Bernadette Mederer, and Christin Kühnert for human cell culture and supporting the bioassays testing. S.K. and G.K. received funding from the German Research Foundation (DFG) grant number GRK 2032/2. Open Access funding was enabled by Project DEAL. The graphical abstract was created in BioRender (https://biorender.com/h18t139).

## Author contributions

Conceptualization: SK, LA, MC; Data curation: SK; Formal Analysis: SK, BE, LH; Funding acquisition: LA, MC; Investigation: SK, KG, BE, LH; Methodology: SK, KG, DD, BE, LH, MC; Project administration: MC; Resources: KG, BE, LA; Supervision: DD, LA, MC; Visualization: SK; Writing – original draft: SK; Writing – review & editing: all authors

