## Supplementary Material for "Sequential exposure to anoxic/oxic conditions leads to biotransformation and detoxification of sitagliptin in urban hyporheic zones"

### Cell counting

Growth of enrichment cultures was assessed by direct cell counting using epifluorescence microscopy. Specifically, an aliquot of 20 µL of the enrichment cultures was mixed with 1.3 µL of SYBR Green I solution (Invitrogen, USA). Then, cells were immobilized by pipetting 18 µL of the stained culture on an agarose-coated glass slide and covering it with a 20 mm × 20 mm cover glass [118]. Subsequently, the stained culture was excised through an epifluorescence microscope (Optiphot 2, Nikon, Minato City, Japan) to take at least 15 distinct micrographs with a Nikon DS-Vi1 digital camera. Micrographs were processed automatically using ImageJ (https://imagej.nih.gov/ij/) to acquire cell counts as described previously [119].

### Analytical techniques

Methane was quantified via direct injection of 250 µL headspace into an 8890 (Agilent, USA) gas chromatograph system equipped with a flame ionization detector (GC-FID) and a GS-Q column (0.5 mm x 30 m, Agilent, USA) as well as with a G1540A (HP 6890 series, Hewlett Packard, USA) gas chromatography system equipped with a thermal conductivity detector (GC-TCD) and a HP 19096C-010 column (PoraPak Q, 80/100 mesh, Hewlett Packard, USA). The system was calibrated with a mixture of $CH_4$/air in defined ratios from 0% to 100% (v/v).

Nitrate and sulfate in microcosms were quantified via a Dionex ICS-5000 ion chromatograph system (Thermo Scientific, Germany) equipped with a Dionex IonPac AS18 column (2 mm × 150 mm, Thermo Scientific, Germany) as described previously [46]. Acetate was quantified using a Dionex ICS-5000+ ion chromatography system (Thermo Scientific, Germany) equipped with a Dionex IonPac AS18 pre- and analytical column (precolumn: 2 mm × 50 mm, analytical: 2 mm × 250 mm, Thermo Scientific, Germany). Potassium hydroxide and milli-Q water were used as the mobile phase with a flow rate of 0.2 mL $min^{-1}$ and the following gradient elution: 0–5 min 10 mM, 5–15 min linear increase from 10 to 40 mM, 15-28 min linear increase from 40 to 60 mM, followed by instant decrease and 28–40 min constant at 10 mM. Detection was achieved through a conductivity detector.

The $pK_a$ value of sitagliptin was measured with a Sirius T3 automated titrator (Pion, Billerica, MA, USA) as described previously [120]. Briefly, 1.15 mg sitagliptin were weighed in a Sirius sample vial. The pH was adjusted to 2 with 0.5 M HCl and the $pK_a$ value was determined by a titration of the sample with 0.5 M KOH. Three titrations were performed from the same sample and the $pK_a$ value reported is the mean of the three replicate titrations.

Pore water samples were analyzed by (Gerundt et al., unpublished manuscript, 2025) as follows: $Fe^{2+}$ was quantified via graphite atom absorption spectrometry (PinAAcle 900Z, Perkin Elmer, Waltham, USA). $NO_3^-$ and $SO_4^{2-}$ were quantified via a 930 Compact IC Flex ion chromatography system equipped with a conductivity detector and a Metrosep A Supp 7 analytical column at 45 °C (4 mm x 250 mm, Metrohm, Switzerland). 3.6 mM $Na_2CO_3$ was used as the mobile phase with a flow rate of 0.8 mL $min^{-1}$. Ion concentrations exceeding the calibration range were extrapolated based on the calibration curves.

**Mammalian cell micronucleus test**

The *in vitro* micronucleus test with A549 cells was conducted based on the OECD 487 guidelines and previous reports [121, 122] with modifications. Specifically, the human lung carcinoma cell line A549 was cultured in DMEM with Glutamax, supplemented with 10% heat-inactivated FBS and 100 U $mL^{-1}$ Penicillin/Streptomycin. The assay was performed in 384-well plates following a 4-day experimental timeline. On the first day, cells were seeded at 750 cells per well and incubated at 37°C in 5% $CO_2$ for 24 h. On the next day, test compounds were introduced, and the cells were incubated for another 24 h. Colchicine served as a positive control. On the third day, cells were washed to remove the test compounds, and again incubated for 24 h (recovery period). On the fourth day, cells were washed again before DNA being stained with Hoechst 33342. Micronucleus (MN) formation was assessed using a high-content microscope (Molecular Devices, San José, CA, USA)) and ImageXpress. Confluency was measured daily with an IncuCyte S3 live-cell imaging system (Essen BioScience, Ann Arbor, MI, USA). The data acquired 24 h and 72 h after seeding were used to derive the growth factor (GF) of the A549 cells:

$$\text{Growth factor (GF)} = \frac{\text{Confluency 72 h [\%]}}{\text{Confluency 24 h [\%]}}$$

The GF of the samples in comparison to the average GF of unexposed cells was used to determine the cell viability:

$$\text{Cell viability} = \frac{\text{GF sample}}{\text{Average GF unexposed cells}}$$

The total number of MN and the total number of cells was measured at 2 sites of each well and the number of MN was divided by the total number of cells to derive the fraction of cells with MN (%MN).

$$\%\text{MN} = \frac{\text{No. of micronuclei}}{\text{Total No. of cells}}$$

To derive the induction ratio (IR) the %MN in the sample was divided by the average %MN in the unexposed cells:

$$\text{IR} = \frac{\%\text{MN sample}}{\text{Average }\%\text{MN unexposed cells}}$$

As no concentration-response model could be applied to the rather variable IR data, the lowest observed effect concentration (LOEC) was defined as the lowest concentration that showed a significant increase in MN induction compared to the unexposed cells. All individual concentrations tested for each chemical that caused less than 50% cytotoxicity were compared with the unexposed cells using a one-way analysis of variance (ANOVA).

### Supplementary Figures

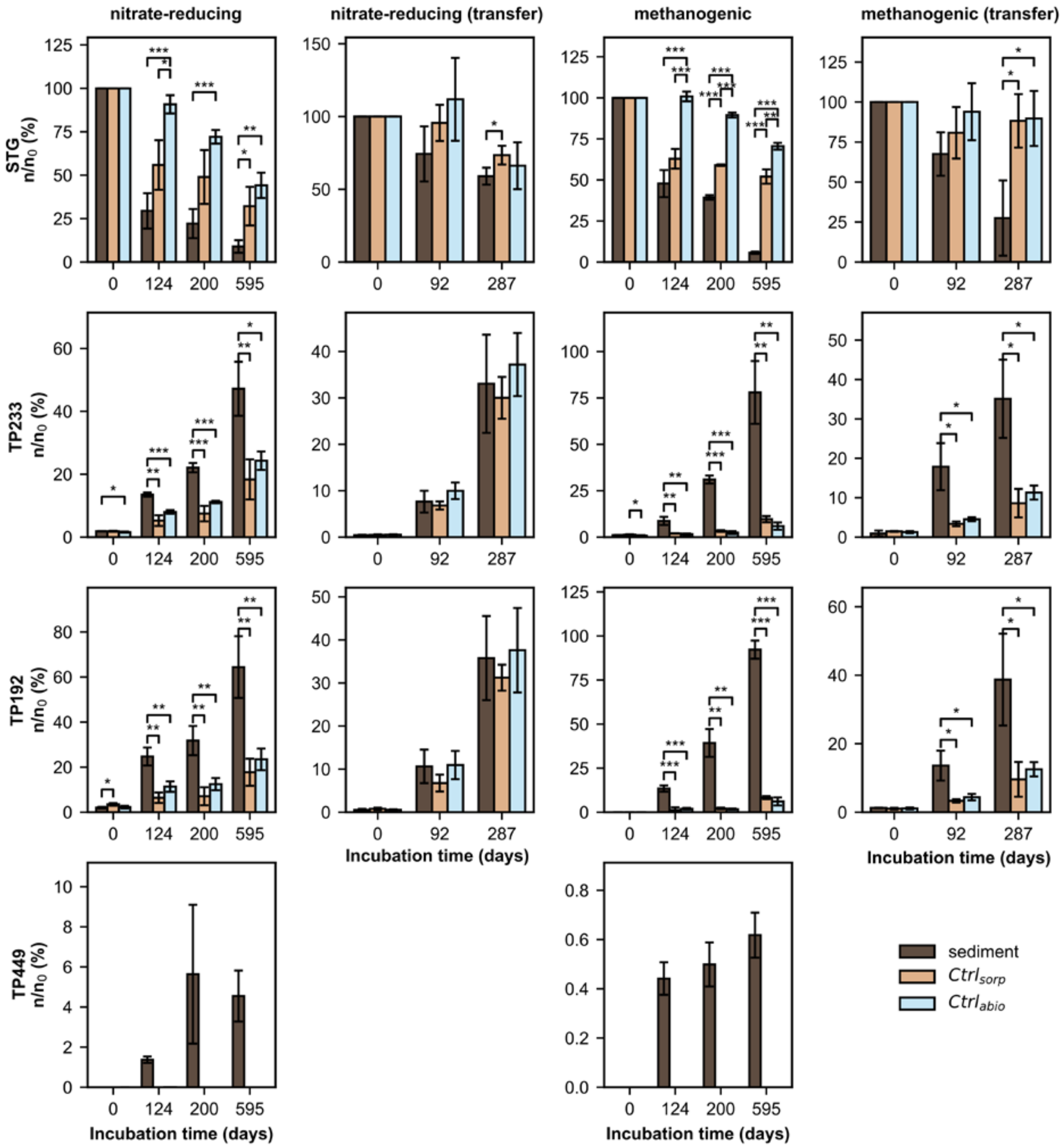


Supplementary Figure 1: Relative molar concentrations of sitagliptin (STG), TP233, TP192, and TP449 in sediment microcosm setups compared to sorption and abiotic controls. Relative concentrations were calculated in relation to the molar STG concentration at day 0. Values are means of biological triplicates. Error bars denote standard deviations. Statistical differences between setups are indicated with '*', '**', '***' based on an independent Student's t-test with $p < 0.05$, $p < 0.005$, and $p < 0.001$, respectively.

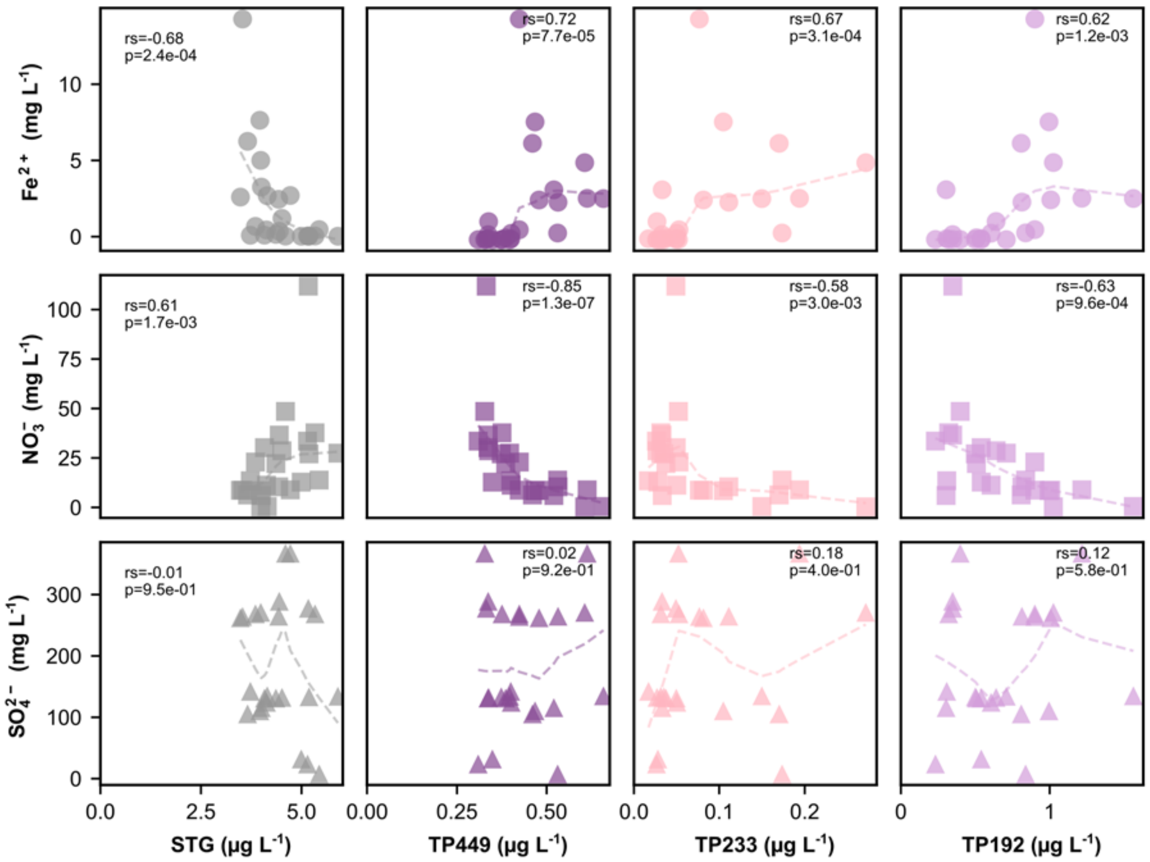


Supplementary Figure 2: Correlation analysis of sitagliptin (STG), TP449, TP233, and TP192 with iron(II) ($Fe^{2+}$), nitrate ($NO_3^-$), and sulfate ($SO_4^{2-}$) concentrations in pore water samples collected from the River Erpe at Site B (sandy). Dashed lines represent trend lines based on locally weighted scatterplot smoothing (LOWESS) with a smoothing factor of 0.7. Spearman's rank correlation coefficients (rs) and corresponding p-values from two-tailed tests are shown in each subplot.

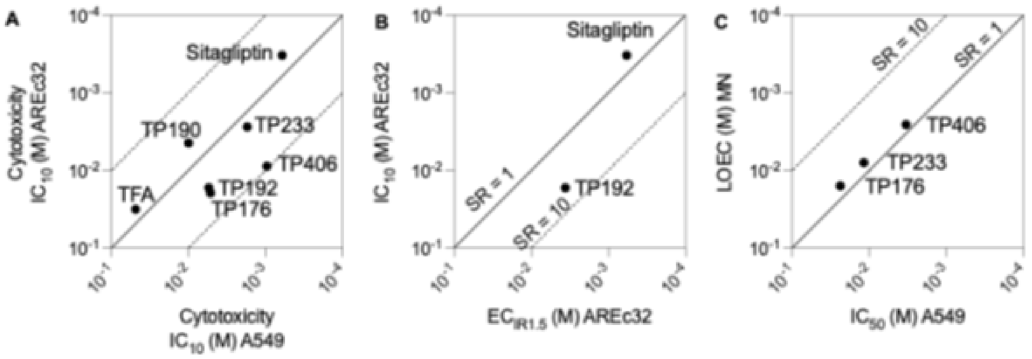


Supplementary Figure 3: Comparison of the effects in the AREc32 and A549 bioassays. A: Comparison of cytotoxicity $IC_{10}$ determined in both bioassays. B: Specificity of the oxidative stress response, specificity ratio SR = $IC_{10}/EC_{IR1.5}$. C: Specificity of the micronucleus formation, specificity ratio SR = $IC_{50}$/LOEC.

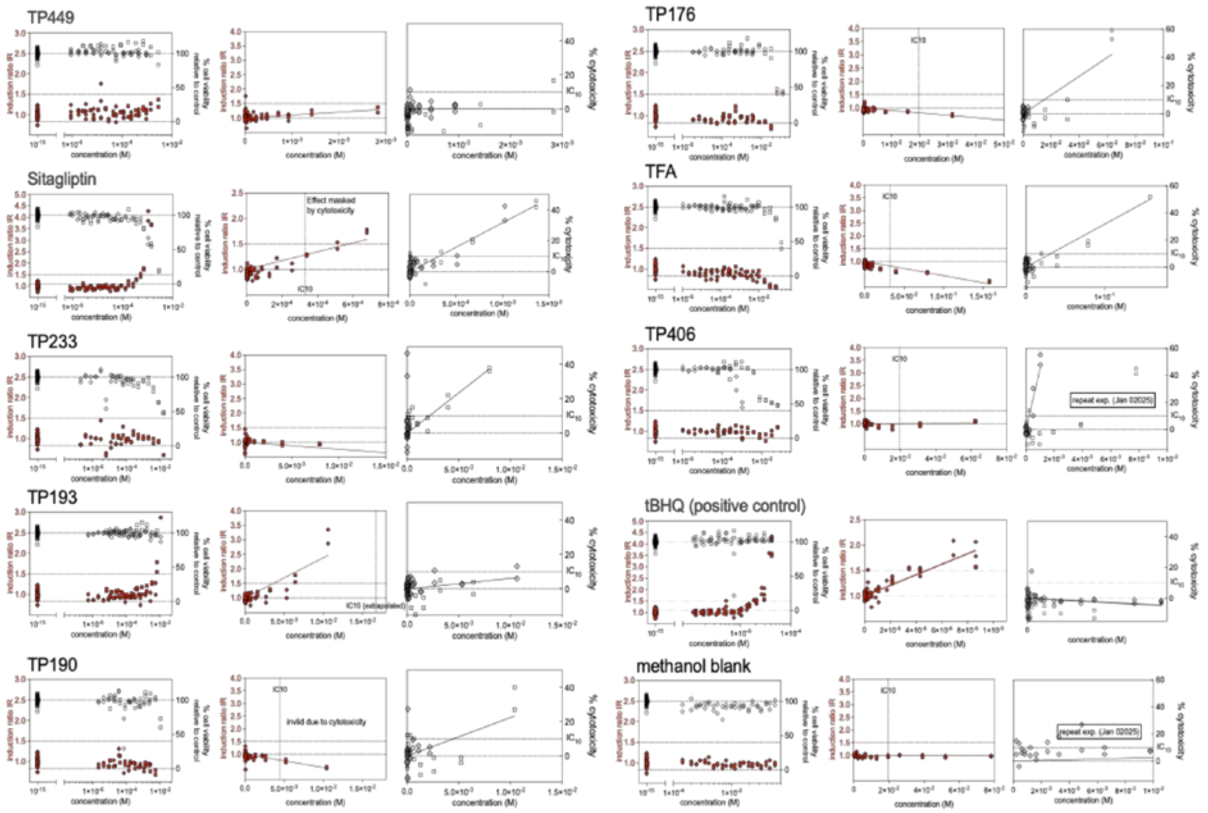


Supplementary Figure 4: Dose-response curves for the AREc32 bioassay.

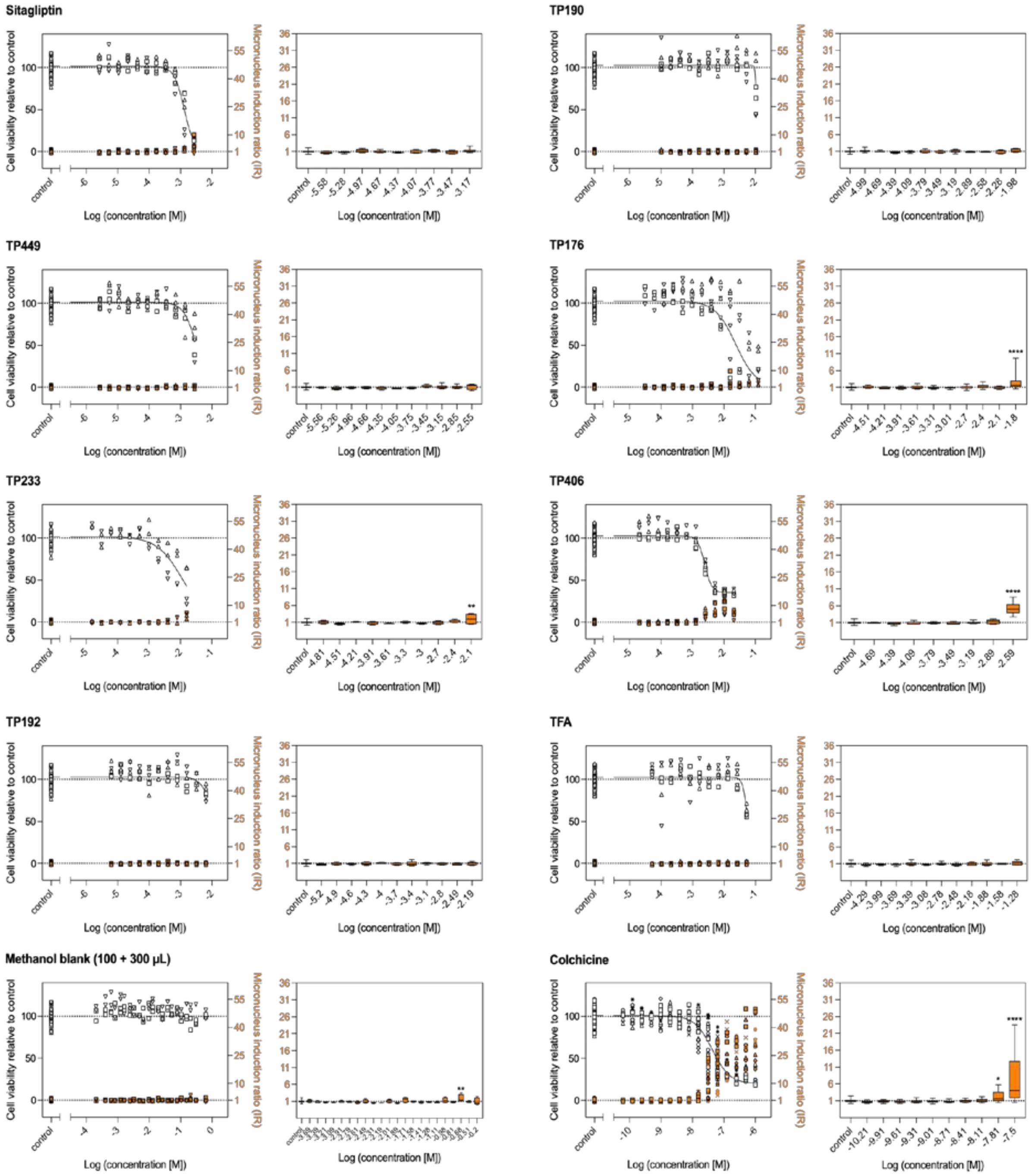


Supplementary Figure 5: Dose-response curves for the micronucleus test.

**Supplementary Tables**

Supplementary Table 1: Gradient used for LC-MS/MS data acquisition in MRM mode.

| Time(min) | A% | B% |
|---|---|---|
| 0 | 95 | 5 |
| 1.5 | 95 | 5 |
| 2.5 | 50 | 50 |
| 9 | 5 | 95 |
| 11 | 5 | 95 |
| 11.1 | 95 | 5 |
| 16 | 95 | 5 |

Supplementary Table 2: Transitions and MRM parameters used for the detection of sitagliptin (STG) and TPs. Transitions used for quantification are marked in bold. DP = declustering potential, CE = collision energy, CXP = cell exit potential. Compounds are marked with ‘x’ if detected in aerobic, nitrate-reducing, or methanogenic microcosms as well as in the River Erpe, respectively. Confidence levels (Lvl) are based on [123].

| **Analyte** | **$R_t$ (min)** | **Ionization mode** | **Precursor (*m/z*)** | **Fragment (*m/z*)** | **DP (V)** | **CE (V)** | **CXP (V)** | **Detected in** | | | | **Lvl** |
|---|---|---|---|---|---|---|---|---|---|---|---|---|
| | | | | | | | | **$O_2$** | **$NO_3^-$** | **$CH_4$** | **Erpe** | |
| STG | 7.07 | + | 408.1 | **235.2** | 45 | 26 | 10 | x | x | x | x | 1 |
| | | | | 193.2 | 45 | 34 | 9 | | | | | |
| | | | | 174.1 | 45 | 35 | 11 | | | | | |
| TP192 | 1.33 | + | 193.1 | **150.1** | 52 | 29 | 24 | - | x | x | x | 1 |
| | | | | 138.2 | 52 | 25 | 29 | | | | | |
| | | | | 118.0 | 52 | 40 | 18 | | | | | |
| TP233 | 6.62 | + | 234.1 | **174.2** | 35 | 23 | 10 | - | x | x | x | 1 |
| | | | | 154.1 | 35 | 38 | 8 | | | | | |
| | | | | 127.0 | 35 | 49 | 12 | | | | | |
| TP449 | 8.57 | + | 450.1 | 193.1 | 43 | 25 | 12 | - | x | x | x | 1 |
| | | | | **174.1** | 43 | 46 | 30 | | | | | |
| | | | | 216.1 | 43 | 32 | 30 | | | | | |
| TP190 | 8.47 | - | 145.0 | **125.0** | -10 | -20 | -10 | x | - | - | x | 1 |
| TP176 | 8.44 | - | 175.0 | 131.0 | -10 | -32 | -10 | x | - | - | x | 1 |
| | | | | **111.0** | -10 | -25 | -10 | | | | | |
| | | | | 91.0 | -10 | -25 | -10 | | | | | |
| TP406 | 8.61 | + | 407.1 | **193.1** | 62 | 34 | 22 | - | - | - | x | 1 |
| | | | | 145.0 | 62 | 58 | 30 | | | | | |
| TFA | 1.72 | - | 113.0 | 113.0 | -38 | -10 | -5 | - | - | - | x | 1 |
| | | | | **69.0** | -38 | -20 | -10 | | | | | |
| TP232 | 1.16 | + | 233.0 | **155.0** | 35 | 38 | 8 | - | - | - | x | 3 |
| | | | | 127.0 | 35 | 49 | 12 | | | | | |
| | | | | 175.0 | 35 | 23 | 10 | | | | | |
| TP435 | 8.37 | + | 436.1 | 244.1 | 60 | 40 | 10 | - | - | - | x | 3 |
| | | | | 408.1 | 60 | 40 | 10 | | | | | |
| | | | | **193.1** | 60 | 40 | 10 | | | | | |
| TP507 | 8.40 | + | 508.1 | 391.1 | 60 | 40 | 10 | - | - | - | x | 3 |
| | | | | 174.1 | 60 | 40 | 10 | | | | | |
| | | | | **193.1** | 60 | 40 | 10 | | | | | |
| Diphenhydramine | 7.86 | + | 256.1 | 167.1 | 10 | 20 | 8 | - | - | - | - | 1 |
| | | | | 152.1 | 10 | 52 | 8 | | | | | |
| SMX-D4 | 6.88 | + | 258.2 | 160.0 | 55 | 22 | 12 | - | - | - | - | 1 |

Supplementary Table 3: Summary of detected amplicon sequencing variants (ASVs) annotated with SILVA 138.2 and GTDB r220 taxonomy. All graphs showing relative taxa abundances from 16S gene amplicon sequencing data are based on this data.

Supplementary Table 4: Nitrate ($NO_3^-$) concentrations in mM in nitrate-reducing microcosm setups determined by ion chromatography and compared to control setups with autoclaved sediment ($Ctrl_{sorp}$) or no sediment ($Ctrl_{abio}$). Values represent means of biological triplicates ± standard deviation.

| **Setup\Incubation time (days)** | **Non-passed** | | | **Passed** | | |
|---|---|---|---|---|---|---|
| | **0** | **124** | **200** | **0** | **92** | **287** |
| Sediment | 0.87 ± 0.03 | 0.01 ± 0.01 | 0.00 ± 0.00 | 0.44 ± 0.10 | 0.01 ± 0.00 | 0.00 ± 0.00 |
| $Ctrl_{sorp}$ | 0.93 ± 0.02 | 0.93 ± 0.05 | 0.93 ± 0.00 | 0.77 ± 0.20 | 0.79 ± 0.20 | 0.82 ± 0.19 |
| $Ctrl_{abio}$ | 0.91 ± 0.04 | 0.95 ± 0.01 | 0.94 ± 0.00 | 0.79 ± 0.02 | 0.79 ± 0.04 | 0.66 ± 0.17 |

Supplementary Table 5: No transformation of sitagliptin observed in sulfate-reducing microcosms shown as dissipation of STG compared to control setups with autoclaved sediment ($Ctrl_{sorp}$) or no sediment ($Ctrl_{abio}$) controls. Microcosms were spiked with 700 nM sitagliptin and about 1 mM $K_2SO_4$ at day 0. Relative abundances were calculated in relation to the molar sitagliptin concentration at day 0. Values represent means of biological triplicates ± standard deviation.

| **Setup** | **Incubation time (days)** | **0** | **318** |
|---|---|---|---|
| Sediment | $SO_4^{2-}$ (mM) | 1.41 ± 0.03 | 0.00 ± 0.00 |
| | Sitagliptin $c/c_0$ | 1.00 ± 0.00 | 0.82 ± 0.07 |
| $Ctrl_{sorp}$ | $SO_4^{2-}$ (mM) | 1.43 ± 0.01 | 1.39 ± 0.08 |
| | Sitagliptin $c/c_0$ | 1.00 ± 0.00 | 0.94 ± 0.10 |
| $Ctrl_{abio}$ | $SO_4^{2-}$ (mM) | 1.43 ± 0.08 | 1.39 ± 0.04 |
| | Sitagliptin $c/c_0$ | 1.00 ± 0.00 | 0.98 ± 0.09 |

Supplementary Table 6: Amount of methane ($CH_4$) in the headspace and acetate in the liquid phase of methanogenic microcosm setups determined by gas and ion chromatography, respectively. $NaHCO_3$ consumption was calculated under the assumption that all methane and acetate originated from $NaHCO_3$ and that no alternative pathways consumed $NaHCO_3$. Values represent means of biological triplicates ± standard deviation. ND = not determined

| **Incubation time (days)** | **0** | **118** | **200** | **494** | **Pass1 (287)** |
|---|---|---|---|---|---|
| Methane (µmol) | 0 ± 0 | 51.3 ± 5.4 | ND. | 208.5 ± 34.7 | 80.2 ± 37.0 |
| Acetate (µmol) | 0 ± 0 | 18.0 ± 14.0 | 33.1 ± 13.9 | 34.8 ± 7.6 | 36.7 ± 6.3 |
| $NaHCO_3$ consumed (%) | 0% | 29.7 ± 10.0 | ND | 101.6 ± 13.9 | 47.2 ± 21.8 |

Supplementary Table 7: Relative sitagliptin concentrations in aerobic microcosms s compared to control setups with autoclaved sediment ($Ctrl_{sorp}$) or no sediment ($Ctrl_{abio}$). Microcosms were spiked with 700 nM sitagliptin at day 0 and incubated in cotton-plugged flasks on a rotary shaker. Relative abundances were calculated in relation to the molar sitagliptin concentration at day 0. Values represent means of biological triplicates ± standard deviation.

| **Setup\Incubation time (days)** | **0** | **36** | **106** |
|---|---|---|---|
| Sediment | 1.00 ± 0.00 | 0.89 ± 0.11 | 0.80 ± 0.09 |
| $Ctrl_{sorp}$ | 1.00 ± 0.00 | 0.82 ± 0.02 | 0.83 ± 0.06 |
| $Ctrl_{abio}$ | 1.00 ± 0.00 | 1.01 ± 0.10 | 0.90 ± 0.05 |

Supplementary Table 8: Cell counts as $10^7$ cells $mL^{-1}$ in methanogenic and nitrate-reducing microcosm setups determined by fluorescence microscopy. Values represent means of biological triplicates ± standard deviation. After 124 days of incubation cell numbers significantly increased in methanogenic samples (Welch's t-test, $p < 0.01$) but stayed constant (Welch's t-test, $p = 0.06$) in nitrate-reducing samples.

| **Setup\Incubation time (days)** | **0** | **124** | **200** |
|---|---|---|---|
| Methanogenic | 1.2 ± 0.2 | 4.3 ± 0.4 | 4.0 ± 0.8 |
| Nitrate-reducing | 1.0 ± 0.0 | 0.7 ± 0.1 | 0.6 ± 0.2 |

Supplementary Table 9: Toxicological assessment of sitagliptin (STG) and its transformation products (TPs) including trifluoroacetic acid (TFA), a putative end product via the *in vitro* AREc32 and micronucleus tests. NE: no effect observed. up to 10% cytotoxicity (AREc32) or 50% cytotoxicity (A549). TR = toxic ratio, SR = specificity ratio, LOEC = lowest observed effect concentration, $IC_{10,baseline}$ = predicted baseline cytotoxicity [124]. Values indicate means ± standard errors.

| **Chemical** | **TP449** | **STG** | **TP406** | **TP233** | **TP192** | **TP190** | **TP176** | **TFA** |
|---|---|---|---|---|---|---|---|---|
| $IC_{10}$ AREc32(mM) | > 2.83 | 0.33 | 8.83 | 2.75 | 16.71 | 4.44 | 19.6 | 31.8 |
| | | ± 0.01 | ± 1.27 | ± 0.30 | ± 6.24 | ± 1.57 | ± 1.55 | ± 1.90 |
| $IC_{10,baseline}$ (mM) | 0.85 | 3.47 | 7.26 | 17.37 | 1.14 | 2.76 | 46.17 | 23.21 |
| TR AREc32 | | 10.55 | 0.82 | 6.31 | 0.07 | 0.62 | 2.35 | 0.73 |
| $EC_{IR1.5}$ AREc32(mM) | > 2.83 | 0.58 | NE | NE | 3.61 | NE | NE | NE |
| | | ± 0.06 | | | ± 0.27 | | | |
| SR AREc32 ($IC_{10}/EC_{IR1.5}$) | - | 0.57 | - | - | 4.63 | - | - | - |
| $IC_{50}$ A549 (mM) | 3.18 | 1.32 | 3.33 | 11.80 | >6.46 | 11.00 | 23.80 | 56.00 |
| LOEC A549 (mM) | NE | NE | 2.57 | 7.94 | NE | NE | 15.80 | NE |
| SR A549 ($IC_{50}$/LOEC) | - | - | 1.29 | 1.48 | - | - | 1.50 | - |

Supplementary Table 10: Summary of detected metagenome-assembled genomes (MAGs).

Supplementary Table 11: Summary of proteins identified in TP233-transforming and control microcosms after 7 days of incubation.

Supplementary Table 12: Relative taxa abundances based on the sum of protein abundances assigned to a metagenome-assembled genome and its taxonomy.
